# Spontaneous cancer regression as inversion of spontaneous cancer progression: Malignant melanoma remission as a case-study towards clinical applicability

**DOI:** 10.64898/2025.12.27.696663

**Authors:** Bindu Kumari, Arushi Misra, Ashish Shrivastava, Kiran Bharat Lokhande, Ashutosh Singh, Prasun K. Roy

## Abstract

The spontaneous cancer regression process (SCR) a rather rare paradoxical natural phenomenon, shows that malignant tumors can episodically undergo complete permanent elimination without treatment, and thus may be used for gaining clues to incisive therapeutic innovations. SCR of malignant breast tumors occurs subclinically across human populations (∼22% rate), according to Scandinavian and Wisconsin Screening Registries (monitoring 0.33million and 2.95million population respectively). SCR occurs in discoid melanomas (∼12% rate). SCR process has high novelty with no toxicity nor recurrence, and duplicating SCR process is indeed clinically desirable. We aim to probe SCR using melanoma as illustrative pilot-study, and identify candidate leads that may mimic SCR on the tumor. We investigated molecular biological of melanoma, finding two reciprocal phases: spontaneous-progression and spontaneous-regression, and showed driver genes for melanoma SCR. Then, we pursued network pharmaco-informatics analysis. We found that spontaneous melanoma eradication shows an unexpected finding, in distinct contrast to the prevalent view that immunological processes and immunotherapy are critical for melanoma regression, than targeting DNA. We found that in contradistinction, it is DNA interference process that is primary route for melanoma regression, while anti-tumor immune activation is only secondarily needed. We observed that targeting two pathways (kinase-system, inositide-system) by inhibiting two genes, arrested melanoma-progression phase, and activated the reverse phase, melanoma-regression. We investigated the interactions of possible candidate leads, indicating significant therapeutic potency. Our findings underscore the high impact possibility for leveraging the anomalous phenomenon of spontaneous cancer regression as a general oncological principle for probing targeted therapeutic interventions, with substantial clinical potentials.

## 1. Introduction

A malignant tumor is usually faced by the physician when it is in the progression phase, nevertheless, the phenomenon of permanent spontaneous regression of malignant tumors is a well-documented biological process [1]. For instance, the spontaneous regression process of malignant breast tumors occurs sub clinically regularly across human populations with a 22– 46% rate, according to Scandinavian and Wisconsin Screening Registries, these registries have monitored a population of 0.33 million and 2.95 million respectively [2,3]. Moreover, it is found from postmortem studies that nearly half of the individuals, in general, have a malignant focus in the prostate or uterus (cervix), with permanent containment, and malignant neuroblastoma tumors are known to fully regress from the original larger-sized tumors [4,5]. The same phenomenology of spontaneous regression occurs in the dermatological malignancy, melanoma. When melanocytes, the pigment cells of the skin, start to grow out without interruption, melanoma tumor formation takes place. Melanoma is very hazardous because, if undiagnosed and untreated, it is considerably more likely to spread to other body regions. Of all cancers, skin cancer is by far the most prevalent. Melanoma is the leading cause of death from the skin cancer, and indeed, the rate of increase of melanoma incidence is one of the highest of all malignancies. Indeed, by the end of the next decade, there will be globally 510,000 new melanoma cases yearly, along with a total 2,050,000 of melanoma patients [6]. Over the past few decades, melanoma rates have been starkly increasing, and efficient treatment of this lethal tumor is urgently required to halt it from growing and spreading elsewhere.

Like many other solid tumors, melanoma may spontaneously regress from an immunological perspective, disappearing completely and permanently [7]. Regression in melanoma is a typical phenomenon to note [8]. It describes the elimination or loss of melanoma that is the result of an immune system reaction against the tumor cells in the host. Regression has been shown in up to 58% of discoid melanomas, where it is a relatively frequent occurrence with an incidence range of 10 to 35% [9]. In fact, an examination of 10,098 patients with melanoma regression revealed that these patients could have strong clinical correlations [10]. Understanding the mechanism behind melanoma regression is crucial for the clinical applicability of the regression mechanism from a therapeutic perspective [11]. Indeed, the episodic natural remission process of malignant melanoma is a model system to investigate the phenomenon of spontaneous cancer regression [12]. Spontaneous cancer regression occurs when a tumor which initially grows spontaneously, undergoes retardation and thence reversion, whereby the tumor gradually undergoes involution and finally eradication. Thus, in the initial phase, there is spontaneous tumor progression which will be actuated by the tumor-promoting genes including oncogenes, which induces cell replication of these proliferated cells, i.e. these genes will have upregulation in this phase. Furthermore, in this phase other genes (e.g. cell stability genes) that inhibit the oncogenes or cell proliferation will be downregulated, thereby leading to the progression of the tumor.

For tumors that later undergo spontaneous eradication, the process of tumor reversion, i.e. conversion from progression to regression behaviour of a tumor is well-documented by cytologists and tumor biologists [13,14]. When tumor reversion occurs, there will be a reversal of upregulation of the cell proliferation genes and the reversal of the cell stability genes becomes vice versa. That is during regression, the oncogenes or cell proliferation-related genes become downregulated, while the oncogene-inhibitory gene (or cell-stability genes or anti-apoptosis genes) become upregulated. Indeed, to observe the difference between the two phases, one can perform an incisive analysis of the gene profile in spontaneous tumor progression phase vis-a-vis spontaneous tumor regression phase. This analysis may have the potential to furnish novel molecular targets with insightful prospects of innovative pharmacological approaches. Hence here we give attention to the spontaneous tumor progression phase and spontaneous regression phase, and thereby obtain more molecular biological and therapeutic insights with mechanistic implications. Indeed, using a systems biology approach, we have earlier shown [15,16] that the following three input entities enable the complete permanent regression and elimination of a malignant tumor: (i) DNA interference factor, ii) immunological antitumor effectors as cytotoxic T-lymphocytes, and (iii) cytokines, as lymphocyte-activators, like interleukin-2 (Figure 1(a)).

**Figure 1:**
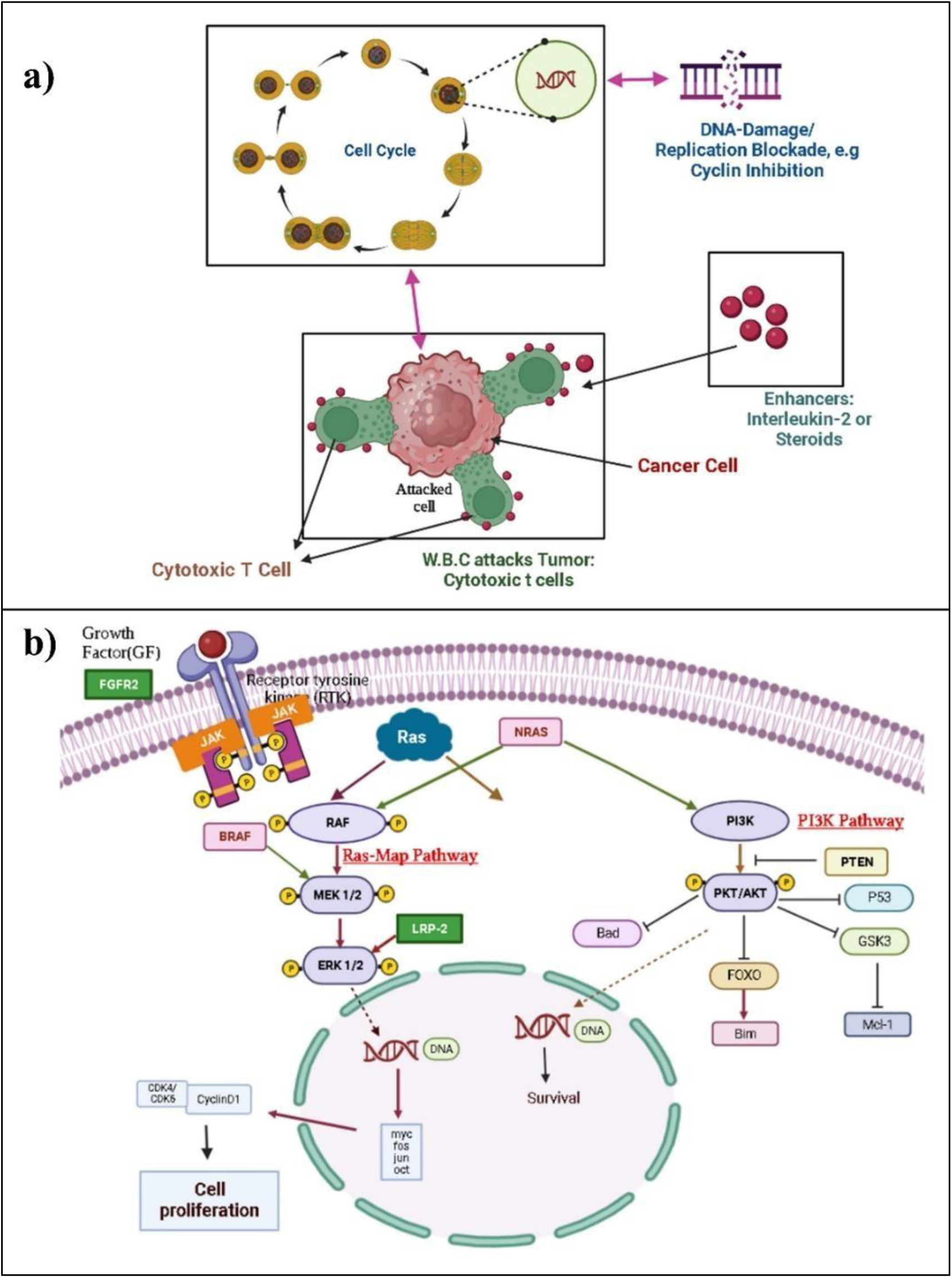
**(a)** Systems biology approach for tumor elimination by three antitumor entities: (i) DNA blockage factor actuating the arrest of malignant cell proliferation, (ii) Immunological antitumor effector as cytotoxic T-lymphocyte and (iii) Lymphocyte activator as Interleukin-2; and (**b**) The formulation of the MAP/ERK and PI3K/AKT signaling pathways in our approach to melanoma proliferation during tumor progression phase, these pathways will downregulate during the spontaneous regression phase.

### 1.1 Tumor progression and regression phases

We here study the melanoma tumor in pigs (Liebechov breed), where the tumor appears at birth and progresses for some time, and then regresses completely, without any relapse, the animals live indefinitely [12]. Initially, after birth, there is tumor growth and a full progression phase across the month, e.g. at t_1_ [i.e., 4 weeks after t_0_, the birth (Schema-1)]. There was requisite animal ethics committee clearance. After time point t_1_, the pigs were assessed at three weekly intervals, namely at time points *t_2_-t_3_-t_4_*, and at the last time point, the melanoma tumor had fully regressed and disappeared. It may be noted that at the second time point t_2_, the tumor progression has halted for an appreciable time, while the regression process has started. The regression process continues to increase through the next time point t_3_; thereafter at the last time point t_4_, the regression process reaches full efficiency producing complete regression of the tumor [12].

**Schema-1:**
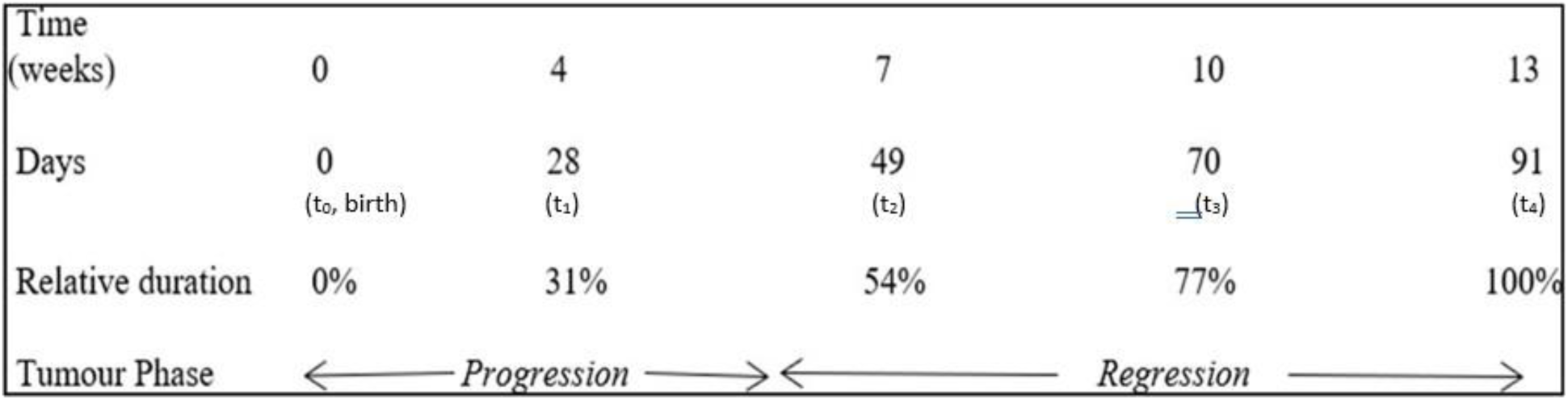
Temporal sequence of the tumor progression phase followed by the tumor regression phase, including complete permanent elimination of the lesion.

Thus, the temporal regime t_1_ pertains to the progression phase, while the temporal regime *t_2_-t_3_-t_4_* pertains to the regression phase, the total duration is about 3 months. In the regression phase, the expression of the genes associated with tumor elimination will have a positive statistically significant value (upregulated) in all three-time points t_2_, t_3_, and t_4_ (if the genes inhibit oncogenes or block cell-proliferation signaling pathways). Correspondingly, if the genes are themselves oncogenes or activate the cell-proliferation signaling pathways, then these genes will be downregulated (statistically significant negative value) during the regression phase. Indeed, the expression of tumor progression genes at t_4_ (when the tumor has been eliminated) will not have statistically significant value. This approach was taken in our previous work [15]. On the other hand, as elucidated earlier, the expression of tumor progression genes at t_1_ will have a statistically significant value.

With respect to the present study (Schema 1), here we deal with a comparison of the maximal progression point (t_1_) and the maximal regression point, i.e. eradication (t_4_). As per the elucidation in the paragraphs above, we may take that the genes (which will be relevant to the tumor reversion process) shall have opposite values of expression at time t_1_ and t_4_, i.e., either upregulation at t_1_ and downregulation at t_4_ or vice-versa.

When considering the likely factors for spontaneous extinction of melanoma, one needs to approach the matter from several aspects, such the gene and pathway factors involved in the regression process, and the possible candidate molecules or drugs that could mimic this process of complete regression on advancing tumors therapeutically. It is desirable also to relate these regression factors to the progression factors, and also have a perspective on the current drugs for melanoma, though often there is tumor relapse, leading to lethality. These factors and drugs, as presently known, are:

*(i). Pathways:* There are a plethora of pathways which are involved melanoma progression, such as Wnt pathway, Notch pathway, MAPK pathway, TGF-β signalling, p53 pathway, PI3K/Akt pathway, INK4 pathway, c-KIT pathway, etc [17].
*(ii). Genes:* There are numerous genes whose mutation are associated with melanoma progression, e.g. NF1, CDK4, KIT, NRAS, BAP1, CDKN2, BRAF, TERT etc [17].
*(iii). Drugs:* Multifarious agents are used as dacarbazine, tomozolomide, fotemustine, trametinib, vemurafenib, and pembroluzimab, in spite of these the 5-year survival of diffuse melanoma is only 22% [17].

Now, shifting the attention to the reciprocal process of spontaneous melanoma regression, the researcher needs to accurately investigate whether the above genes or pathways are downregulated in melanoma regression; this matter is particularly pertinent as one knows that the genes related to the spontaneous regression process *may be different* from the genes associated with spontaneous progression of that disease.

To exemplify the last point, it is known in multiple sclerosis that CD58 gene is associated with regression or remission, while TGFβ1 with progression or relapse [18]. On a similar vein, it transpires that in chronic myelogenous leukaemia, FPGT and TSTA genes are pertinent to remission, while BATF and RAC2 genes with progression [19]. In view of the aforesaid, we need know whether the genes/pathways related to melanoma regression are same or different from the genes/pathways pertinent of melanoma progression. Even if the gene/pathway sets are same, one requires to know which ones of the *regression* genes/pathways are critically important, given that a large of genes may be associated with melanoma *progression* [item (ii) above]. Likewise, the unique pharmaceutical agents that may come out from our spontaneous melanoma regression study may be different from the customary antimelanoma drugs enumerated in item (iii) above.

### 1.2 Reciprocity between progression and regression process

In this study, we analyze the signaling pathways involved in the progression phase of melanoma. Then, we also consider the pathways involved in the reciprocal process, the regression phase of melanoma. In recent years, much progress has been made in understanding the regulatory and genomic processes involved in melanoma initiation and progression. As mentioned above, in our previous work [15], we investigated the regression genes involved in the spontaneous regression phenomenon of melanoma by considering the genes whose expression was consistent from t_2_ to t. (Schema-1) While analyzing that approach, we found that in spontaneous regression, the Top2A gene is key gene which is downregulated. Therein, we had targeted the Top2A gene during melanoma growth and found the candidate drugs (as teniposide or epirubicin) to inhibit this gene, so as to replicate the tumor regression phenomenon. However, in the present study, we have taken a complementary approach to find out the genes that may have effect on melanoma elimination. These genes we will now consider, and accordingly we take those genes whose expression levels differ between time points t_1_ (progression) and t_4_ (regression), and these genes could be pharmacologically targeted for enabling tumor extinction.

To have a perspective on the melanoma progression process, the genetic alteration in mitogen-activated protein kinase (MAPK) and the phosphoinositide 3-kinase (PI3K) signaling pathways form a main factor profile actuating melanoma progression. MAPK and PI3K signaling pathways help melanoma in proliferation, invasion, survival, and angiogenesis [20]. In fig. 1 we show how both pathways are activated by Ras protein mutation (**Figure 1 (b)**). The v-Raf murine sarcoma viral oncogene homolog B (BRAF) is a crucial mutated oncogene in melanoma, with the BRAF V600E mutation being a most significant mutation, accounting for 40–50% of all mutated melanomas and 80% of BRAF-mutated tumors [21]. The neuroblastoma RAS viral oncogene homolog (NRAS), which is mutated in over 30% of all melanomas, is the second most significant mutant oncogene [21].

In this present report, we systemically probe the signal pathways driving melanoma progression and regression. Regarding the differentially expressed genes, we performed the analysis of the molecular function, biological processes, and cellular component. We investigated the tumor elimination process with respect to Fig. 1(a), particularly the actuating inputs: (i) DNA interference factors, and the Immunological antitumor factors, as (ii) immune effector cells and (iii) cytokines. Moreover, we investigated the main driver genes and the associated pathways and targets, and the possible anti-melanoma drugs thereby implied. To substantiate the feasibility of the pharmacological agents identified, we performed the docking study and molecular dynamic analysis to substantiate our approach that the drugs that would thereby mimic or replicate the spontaneous tumor regression process on melanoma lesion of patients, without the risk of tumor recurrence, cancer stem cell relapse, or systemic toxicity, the well-known hallmarks of the spontaneous remission.

## 2. Materials and Methods

### 2.1 Systems biology approach to spontaneous cancer regression process

Whether it is an exogenous tumor remission (induced by therapeutic agents) or an endogenous tumor remission process (host tissue-initiated), the remission process may be described by first-order reaction rate kinetics for tumor cell lysis [22]. As a result, the malignant tumor cell population M has an exponentially declining trajectory, which means a small number of tumor cells always remains at the end of the remission process under the asymptotic tail of the curve **(Figure 2(a))**. This small residue of malignant cells multiplies later and is responsible for tumor relapse subsequently.

**Figure 2:**
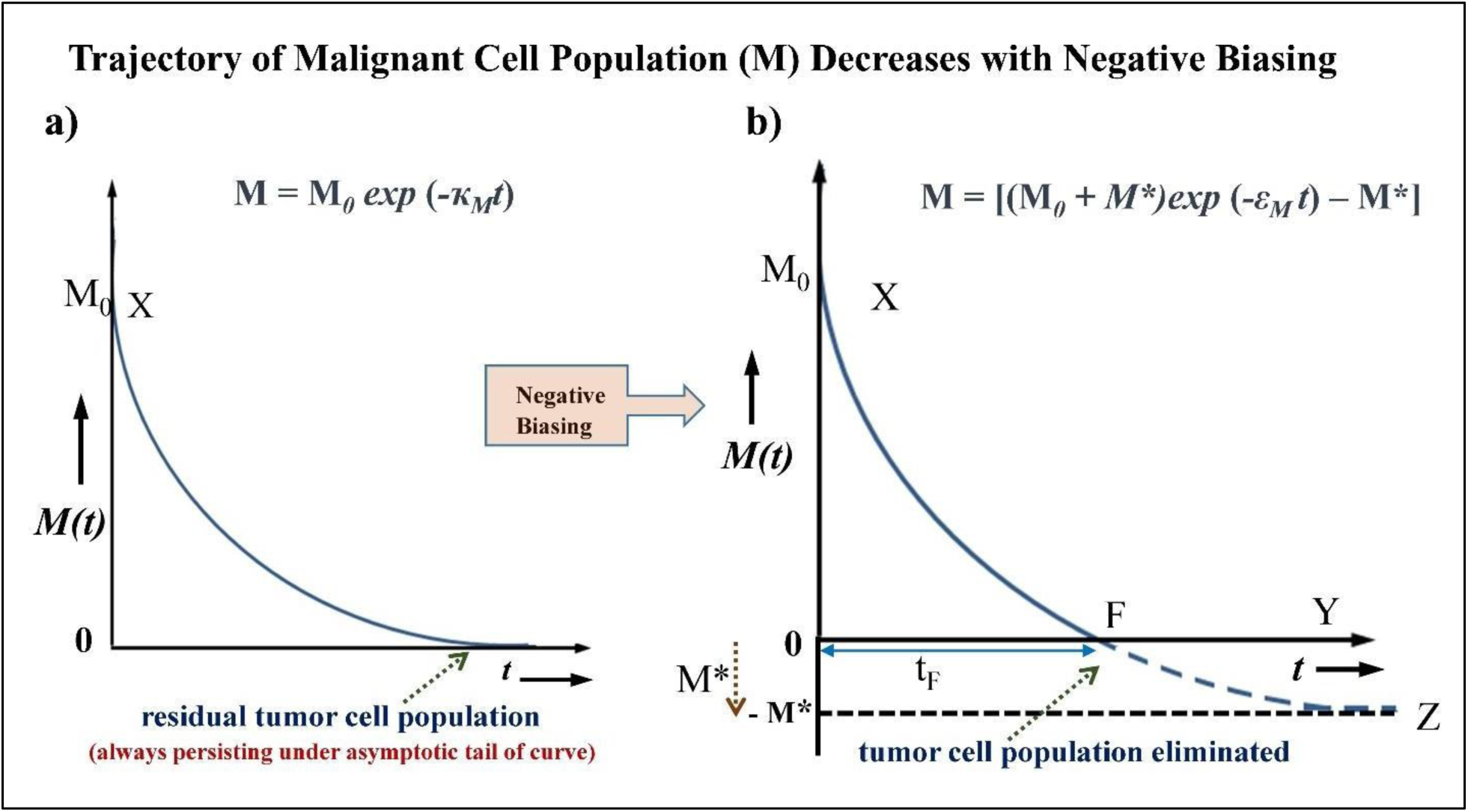
Complete permanent elimination of malignant cell population in a tumor: **(a)** The elimination of the population of tumor cells M(t) follows an exponentially decreasing curve in conventional therapy, there is the persistence of residual tumor cells asymptotically under the tail of the curve, these cells multiply later, and thus produce tumor recurrence.. **(b)** A small negative bias (– M*) enables the exponentially decreasing tumor cell population to become zero at the definitive time point t_F_.

The aforesaid is the usual process of tumor remission, and the majority of tumors that are therapeutically treated follow this trajectory. However, in the case of permanent spontaneous cancer regression, the tumor cell population is eliminated completely and for all future time. To account for this process, we have introduced the concept of the negative biasing technique in our earlier research, by which tumor cell trajectory shifts towards a negative value M*, and thereby all malignant residual cell population become zero at definite time point t_F_ **(Figure 2(b))**; this approach is also delineated in our earlier work [15,16].

It may be mentioned that regarding our model of exponential decrease of malignant cell population during spontaneous tumor regression and eradication, has collateral verification from the framework of systemic homeostasis and cell damage control. To clarify, it has been shown that spontaneous tumor regression process will definitively display exponentially declining behaviour, this exponential dynamics occurs since only this form of the curve displays optimization of the trajectory, so that the total systemic damage toxicity cost to the organism is minimum [23]. As shown therein, the total systemic cost is estimated by the amount of malignant tissue load (neoplastic cell population) eliminated, as characterized by the cumulative neoplastic cell damage or hits, across the time duration of the regression process. This minimization behaviour is expression of the generalized principle of least energy cost observed in biological self-reparative processes [24].

We validated our negative bias formulation **(Figure 2(b)**) using experimental findings on permanent spontaneous regression of various malignancies, such as fibrosarcoma, histiocytoma, and, melanoma tumors [15,16]. Furthermore, to enable this regression, we experimentally found the level of small bias M* to be only about 1% of the initial tumor load M_0_ [15]. In our aforesaid works, utilizing a systems biology approach, we formulated a multimodal equivalence of two permanent tumor regression processes, whether endogenous or exogenous, as induced respectively by host tissue environment or oncological therapy. As we have delineated before [15,16], these two regression processes are actuated by three input factors: (i) DNA blockage factor, (ii) Cytotoxic T-cells, and (3) Interleukin-2 (**Figure 1 (a)**). Additionally, six different differential equations were formulated to depict the interaction between these three input elements and the tumor cell population [15,16]. The time-wise orchestration of these three anti-tumor entities occurs by actuation of various genes and signal transduction pathways. We here try to delineate these genes and pathways by systematic analysis of the microarray findings from melanoma tumors as they underwent spontaneous progression initially, followed by spontaneous regression.

### 2.2 Investigation of spontaneous progression and regression process in malignant neoplasm

In our earlier study, we had analyzed the raw microarray preparation of the spontaneous tumor regression behaviour of melanoblastoma-bearing Libechov minipigs (MeLiM) from the ArrayExpress platform (https://www.ebi.ac.uk/biostudies/arrayexpress/studies/E-MEXP-1152?query=E-%20MEXP-1152) using R-studio platform [15]. Here, the tumor spreads and kills about half of the pigs. Nevertheless, in the other half of animals, the tumor develops up to a certain point (tumor progression), and then undergo spontaneous shrinking and healing (tumor regression), and these latter animals remain healthy thereafter.

### 2.3 Identification of differentially expressed genes (DEGs) of spontaneous melanoma progression and regression regression

After applying the two primary cutoff criteria (−2 >FC > +2 and p-value < 0.05), we have obtained 70, 322, 1147, and 1349 differentially expressed genes (DEGs) at the above-mentioned time points t_1_, t_2_, t_3_, and t_4_. For this current study, we have taken two types of approaches:

a. Type-1: As in our prior study, we had described/considered how the regression process started from time point t_2_; hence we only included genes whose expression was consistent across the regression process i.e. from t_2_ to t_4_.
b. Type-2: The other gene sets that we consider now are those genes whose expression levels differ between time points t_1_ (full progression) and t_4_ (full regression); for example, a gene that is upregulated at t_1_ should be downregulated at t_4_, and vice-versa.

We also consider the protein-protein interaction (PPI) networks which are protein complexes arranged in networks that are consolidated by electrostatic or biochemical factors [25]. The PPI network is essential for molecular functions, and we use the String platform to generate the protein-protein interaction networks for selected DEGs.

### 2.4 Gene Ontology Analysis

The Gene enrichment analysis of DEGs is performed by the DAVID platform [26] (https://david.ncifcrf.gov/), FunRich facility (version: 3.1.3) [27] and ClueG0-v2.5.5 platform from Cytoscape [28]. Using the Gene Ontology (GO) approach and the p-value, we then graded the significant terms for (i) Cellular component (CC), (ii) Biological process (BP), and (iii) Molecular function (MF). As a result, in this study, we created and compared the network of functionally related GO term using the standard adjusted kappa statistic of ҡ > 0.4. FunRich analysis was employed in the present study to examine the biological pathways involved in DEGs. The top 5 biological pathways for genes that were upregulated and downregulated, respectively, were then displayed as bar charts. Statistics are considered significant at p<0.05.

### 2.5 Investigation of the genes

Now, we need to identify the most effective melanoma-regressing driver gene profile from the aforementioned two types of approaches for selecting DEGs. For the first case, we have used the Cytohubba plugin from Cytoscape, already explained in our previous work [15]. For the second case we have selected the common gene among:

i. the pig spontaneous melanoma microarray data genes [i.e. those genes whose expression levels have opposite signs (negative or positive) at time points t_1_ and t_4_], and
ii. the human melanoma gene profile from NCBI database and the human melanoma gene profile from Gene card database.

### 2.6 Identification of target proteins

Targeted molecular therapies have mostly been recognized as a novel therapeutic procedure in most malignancy types. We may target some of the signaling pathways involved in malignant melanoma progression pathways of fig. 1(b). For this purpose, small-molecule inhibitors play a crucial role. Recent evidence shows that tumor growth cannot be blocked or stopped by only inhibiting a single effector protein of the signal transduction cascades that play a role in melanoma pathogenesis; however, a multimodal molecular modification and inhibition is necessary for the most effective targeted melanoma treatment [29]. We elucidate the latter aspect below.

#### (i) Melanoma signaling pathway

Recent developments in molecular oncology have produced new treatment approaches that specifically target (i) immune regulatory molecules involved in suppressing the antitumor immune response, such as T-lymphocyte-associated antigen 4 (CTLA4), programmed cell death 1 receptor (PD-1), and its ligand (PD-L1), and (ii) key effectors of the pathways, for instance mutations in BRAF, NRAS or cKIT genes, which are found to play an important role in the pathogenesis of melanoma [30]. We kept these aspects in our attention as we formulated the anti-tumor approach.

#### (ii) MAP Kinase-ERK Dependent Pathway

The aforementioned NRAS and BRAF proteins are part of the mitogen-activated protein kinase (MAPK) signal transduction system, which controls cell growth, survival, and proliferation, and mediates the response of cells to mitotic external stimuli. Three tissue-specific isoforms of the small proteins produced by the RAS gene family—HRAS, KRAS, and NRAS—are linked to the cytoplasmic membrane. Indeed, most NRAS mutations have been found in melanomas [31,32]. Further, RAF and phosphatidylinositol 3 kinase (PI3K) are two downstream cytoplasmic proteins that can be activated by NRAS. The three members of the RAF kinase family—ARAF, BRAF, and CRAF—are proteins whose activation depends on the formation of complexes by these various isoforms [33]. All these three proteins contribute to the signal transduction of MAPK pathway, and BRAF causes the activation of MEK kinase in melanocytes, which then triggers the activation of ERK, the last enzyme in the MAPK cascade (**Figure 1 (b)**). In 40–60% of melanoma cases, the BRAF gene is mutated; the most frequent alteration (occurring in around 90% of cases) is represented by the substitution of valine for glutamic acid at codon 600 (BRAFV600E) [34]. The BRAFV600E variation provides ongoing stimulation of cell proliferation and tumor formation through activating the phosphorylation of ERK, as do the remaining mutations in the BRAF kinase domain. However, the finding that BRAF is even mutated in common nevi implies that melanoma development requires the activation of BRAF’s oncogenic pathway but is not a prerequisite for it [35]. We note these aspects and give focus to them as our analysis develops.

#### (iii) PI3Kinase-AKT Dependent Pathway

The second route that uses RAS to regulate cell proliferation is made up of the signal transduction pathway: PTEN-PI3K-AKT [30]. The activation of PIK3 and the activity of the phosphatase PTEN protein both affect intracellular levels of PIP2 and PIP3 phosphoinositols in physiological settings [36]. High PIP3 levels modulate the synthesis of proteins essential in cell growth and survival as well as apoptosis, by progressively activating downstream AKT (mostly AKT3 in melanoma) and its substrate mTOR. The activation of AKT leads to inhibition of apoptosis by inactivating numerous pro-apoptotic proteins, such as BAD (BCL-2 antagonist of cell death) and MDM2 (which results in the degradation of p53), as well as promoting cell proliferation through the induction and stabilization of CCND1 and through the inhibition of apoptosis [25]. In view of the above, an angle of our approach can be formulated in Figure 1 (b). In conclusion, PTEN inactivation and PI3K-AKT stimulation result in the abnormal proliferation of cancer cells and the development of apoptosis resistance. Around 70% of all melanomas have dysregulated activation of the AKT pathway, which is due to AKT3 amplification and PTEN loss due to epigenetic silencing or deletion, as originally described [37,38]. We consider the dynamics of this aforesaid analysis, as we elucidate the formulation of investigation duly.

### 2.7 Identification of Putative Candidate Pharmacological Agents

Here with the help of DGIdb (The Drug Gene Interaction) platform (https://www.dgidb.org/), we have selected only the FDA-approved molecules, which are targeting both the signaling pathways i.e. BRAF and NRAS. Furthermore, we have taken those molecules whose effects are common in both pathways, and we selected those drugs that have neither been investigated nor used for melanoma. Also, we consider those molecules that are used to target both the pathways in other disorders but have not been studied for melanoma.

### 2.8 Validation of Candidate Drugs using Computational Biology

#### Protein and Antibody Structure Retrieval and Preparation for Molecular Docking

The 3D structures of the proteins were obtained from the Protein Data Bank (PDB) platform. The PDB ID for BRAF is 5C9C [35], with a resolution of 2.70 Å, and for NRAS, it is 6ZIZ [36], with a resolution of 1.78 Å. The selection of the 6ZIZ structure for NRAS was based on the presence of the Q61R mutation, a well-known mutation in melanoma. Similarly, the choice of the 5C9C structure for BRAF was motivated by the inclusion of the V600E mutation, another critical mutation observed in melanoma patients. The structure of Cetuximab, a monoclonal antibody, was obtained from the PDB with the ID 1YY8 [37]. Protein preparation followed the guidelines outlined in the Discovery Studio manual, involving several steps. Initially, the protein underwent cleaning, which entailed the removal of heteroatoms and water molecules. Subsequently, polar hydrogens were added, and the binding site was defined based on the current selection option. The prepared protein was then saved in ‘.pdb’ format, rendering it suitable for docking studies

#### Ligand Structure Retrieval and Energy Minimization

The 3D structures of the drugs thus obtained, from PubChem database in ‘.sdf’ format, are Alpelisib (PubChem CID: 56649450) and Obatoclax (PubChem CID: 11404337). The ligand structure was gathered from PubChem (https://pubchem.ncbi.nlm.nih.gov/) in ‘.sdf’ format and ligand preparation (e.g removal of charges, ionization method which is pH-based etc) was also performed using Discovery Studio.

#### Binding Site Definition

For the NRAS oncogene, the 6ZIZ protein structure was found to be bound to GTP, a Mg2+ ion, and a compound EZZ (3∼{S})-3-[2-[(dimethylamino)methyl]- 1∼{H}-indol-3-yl]-5-oxidanyl-2,3-dihydroisoindol-1-one). To facilitate docking calculations, these ligands were removed, and the GTP binding sites were utilized. Similarly, in the case of the BRAF gene, analysis revealed that the 5C9C protein structure was complexed with a Cl-1 ion and a compound LY3009120 or 4Z5 [(1-(3,3-dimethylbutyl)-3-{2-fluoro-4-methyl-5-[7-methyl-2-(methylamino) pyrido[2,3-d]pyrimidin-6-yl]phenyl}urea]. These compounds were removed, and the binding site for 4Z5 was designated for docking studies. All other parameters were maintained at their default settings.

#### Protein-Ligand Interaction

We enabled the protein-ligand docking by using the LibDock tool under the receptor-ligand interaction menu on Discovery Studio [38]. The algorithm aligns the ligand conformation to hotspots (polar and apolar) of receptor interaction sites and retains the best scoring poses of ligands [39]. Further, CDOCKER, a grid-based molecular docking program that makes use of the CHARMm force field, was used to form a stable complex inside the active site of protein [40]. Ligands with the lowest CDOCKER energy and highest LibDock scores were shortlisted for further Molecular dynamic simulations.

#### Macromolecular (Protein-Antibody) Analysis

Under the Macromolecules section of this platform, we used Dock and Analyze Protein Complexes and docked the Receptor proteins (NRAS and BRAF) with the monoclonal antibody (Cetuximab) and further processed the poses. The Z-Dock protocol provided rigid body docking of two protein structures using the ZDock algorithm [41] as well as clustering the poses according to the ligand position. Processing poses allowed the selection of a subset from a set of docked protein poses generated by the ZDock protocol. After this, we refined the docked poses and finally analyzed the interactions.

Proposed ligands were subjected to molecular docking in order to find the hits that were biologically active. The conformation with the lowest binding energy was taken to form a stable complex inside the active site of NRAS and BRAF. The CDOCKER program was used to carry out this investigation. A greater value denotes a more advantageous binding, and the CDOCKER score is expressed as a negative value (i.e., -CDOCKER_ENERGY). The H-bonds, van der Waals forces, and electrostatic interactions between the target protein and the ligand were used to compute the CDOCKER energy [42]. Based on the crystal data of the template protein, the binding site of the modeled protein was fixed. To find the local minima (lowest energy conformation) of the modeled NRAS and BRAF with an energy gradient of 0.1 kcal mol−1 Å−1, the smart minimizer technique was used in conjunction with the CHARMm force field.

### 2.9 Molecular Structural Dynamic Analysis

We then utilized the Desmond platform for molecular dynamic (MD) simulation study to analyze protein-ligand interaction stability. The MD simulation systems were built using the system builder module of Desmond. Solvation of the system was carried out using the TIP3P water model. Periodic boundary conditions for each system were established using the buffer method and orthorhombic box shape with a 10 Å distance. Counter ions were added to neutralize the system as required. The systems used for the MD simulations were prepared using an OPLS-2005 force field [43]. The MD simulations were performed for 100 ns for each system. The energy was recorded at every 1.2 ps, and the trajectory was saved at every 100 ps. The volume of the dynamical box was equilibrated with the NPT (constant-temperature, constant-pressure) ensemble at 300 K and 1.01325 bar pressure. Before selecting the simulation option, the systems were relaxed using Desmond’s default relaxation protocol, i.e. Relax model system. All MD simulation trajectory analyses were carried out using Desmond’s simulation event analysis tool, which was used to calculate the various relevant properties and information [44].

The Root Mean Square Deviation (RMSD) parameter provides information on a protein’s structural features during the simulation. If the simulation has reached equilibrium, it can be revealed by this parameter. On the other hand, the root mean square fluctuation (RMSF) is calculated to describe the local alterations or variations near the amino acids in the protein chain. Furthermore, the radius of gyration (Rg) reveals the compactness of the protein and its folding.

Since the intermolecular interactions between the protein-ligand complex are crucial measurements to infer the binding stability of the complex, we measured the various types of intermolecular interactions (Hydrogen bonds, salt bridges, and pi interactions). The pertinent figures were generated using Schrodinger’s Maestro.

### 2.10 Binding Free Energy Calculations

The Prime [45] MM/GBSA (Molecular Mechanics/Generalized Born Surface Area) binding free energy calculations were conducted to assess the protein-ligand complexes over a 100 ns MD simulation period. The Prime MM/GBSA method was employed to compute the binding free energies of the protein-ligand complexes. This approach involves dissecting the total binding free energy into various contributing factors, including electrostatic, van der Waals, solvation, and other energetic terms, providing a detailed understanding of the molecular interactions. The binding free energy, assessed through MM/GBSA calculations, was determined using the Schrödinger’s Prime module. This analysis concentrated on the top 10 poses selected from a pool of 1000 frames/trajectories obtained through 100 ns of MD simulation. The equations utilized to compute the binding free energy resulting from ligand binding to Galectins are outlined as follows [46]:

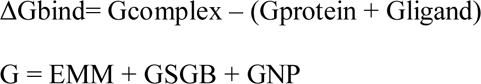

where Gcomplex signifies the energy of the Protein-Ligand complex, Gprotein denotes the energy of the protein, and Gligand represents the energy of the unbound ligand. Additionally, EMM stands for molecular mechanics energies, GSGB represents a SGB solvation model accounting for polar solvation, and GNP indicates the nonpolar solvation term.

The binding free energies obtained, comprising the total binding free energy (ΔBind_Total) and individual contributions from different energy components (such as ΔBind_Coulomb and ΔBind_vdW), were analyzed for each protein-ligand complex. Statistical analyses, including the calculation of mean values and standard deviations, were performed to characterize the variability and reliability of the results [47].

## 3. Results

From the perspective of the natural phenomenon of spontaneous regression of cancer, here we investigated the driving genes and downstream-regulated pathways for replicating this extinction phenomenon in melanoma. Moreover, we have also identified the potential drugs that could block the melanoma progression pathway and thereby enable the elimination of malignant melanoma. Finally, to validate our findings, the molecular docking and molecular dynamic simulation were carried out on the targets and candidate molecules of putative pharmacological agents.

### 3.1 Microarray Data Analysis and Identification of Differentially Expressed Genes

The differentially expressed genes (DEGs) were identified based on fold change value (−2>FC > +2) and p-value (p-value < 0.05). We have taken two types of approaches to investigate the spontaneous regression driving genes as discussed in method section 2.3. From the first approach, we have 176 upregulated and 116 downregulated DEGs whose expression values were consistent from time point t_2_ to t_4_. Likewise, from the second approach, we have 213 DEGs whose expression values have differing signs at time points t_1_ and t_4_. After combining both the approaches and removing the common genes we have obtained 344 upregulated and 160 downregulated DEGs. **Supplementary Figure S1** provides the string network for these DEGs that are respectively upregulated and downregulated.

### 3.2 Gene Enrichment Analysis of Differentially Expressed Genes

With p < 0.05 used as the cutoff, the gene ontology (GO) and pathway enrichment analysis were examined using multiple platforms, including DAVID [26], KEGG pathway [48], and FunRich facility [27]. The DEGs were divided into three functional groups by GO analysis: The Molecular Function (MF), Biological Process (BP), and, Cellular Component (CC) groups as follows.

#### (i) Molecular Function Group

*Upregulation:* For Molecular function (MF), the upregulated genes were related to immunological defense functionality, such as the cytokine binding, and chemokine receptor binding **(Figure 3(a))**.

**Figure 3.**
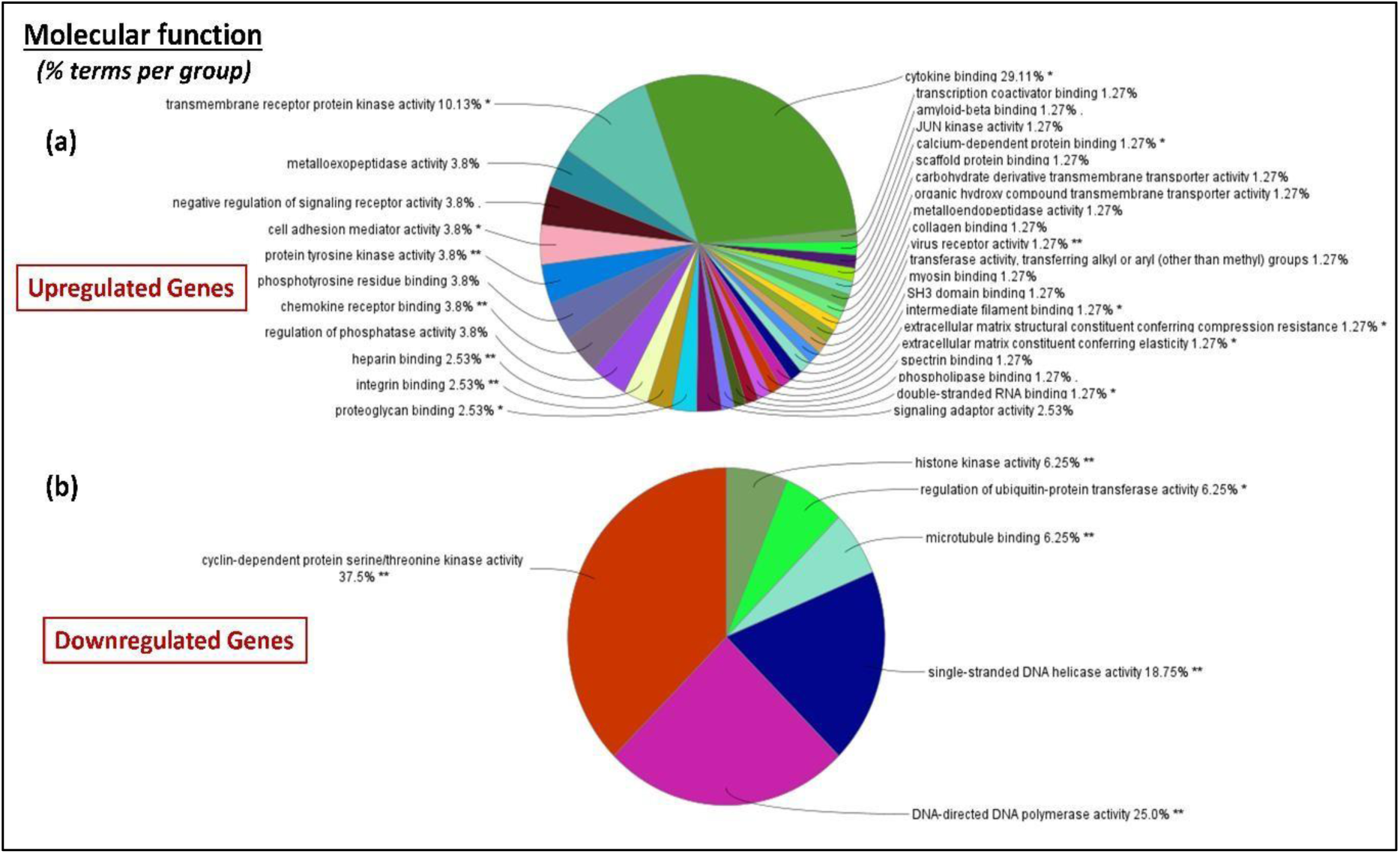
Molecular Function (MF) of enrichment analysis: (a) Upregulated genes; (b) Downregulated genes.

*Downregulation:* However, the downregulated genes were mainly associated with cyclin-dependent protein serine/threonine kinase activity, histone kinase activity, microtubule binding, single-stranded DNA helicase activity and DNA-directed DNA polymerase activity **(Figure 3(b))**.

#### ***(ii)*** Biological Process Group

*Upregulation:* The genes upregulated in the Biological Process (BP) group were enriched in the immune regulation domain, for instance, immune response, activation of immune response, interleukin-27-mediated signalling pathway, antigen receptor-mediated signaling pathway, and leukocyte activation involved in immune response **(Figure 4 (a))**.

**Figure 4.**
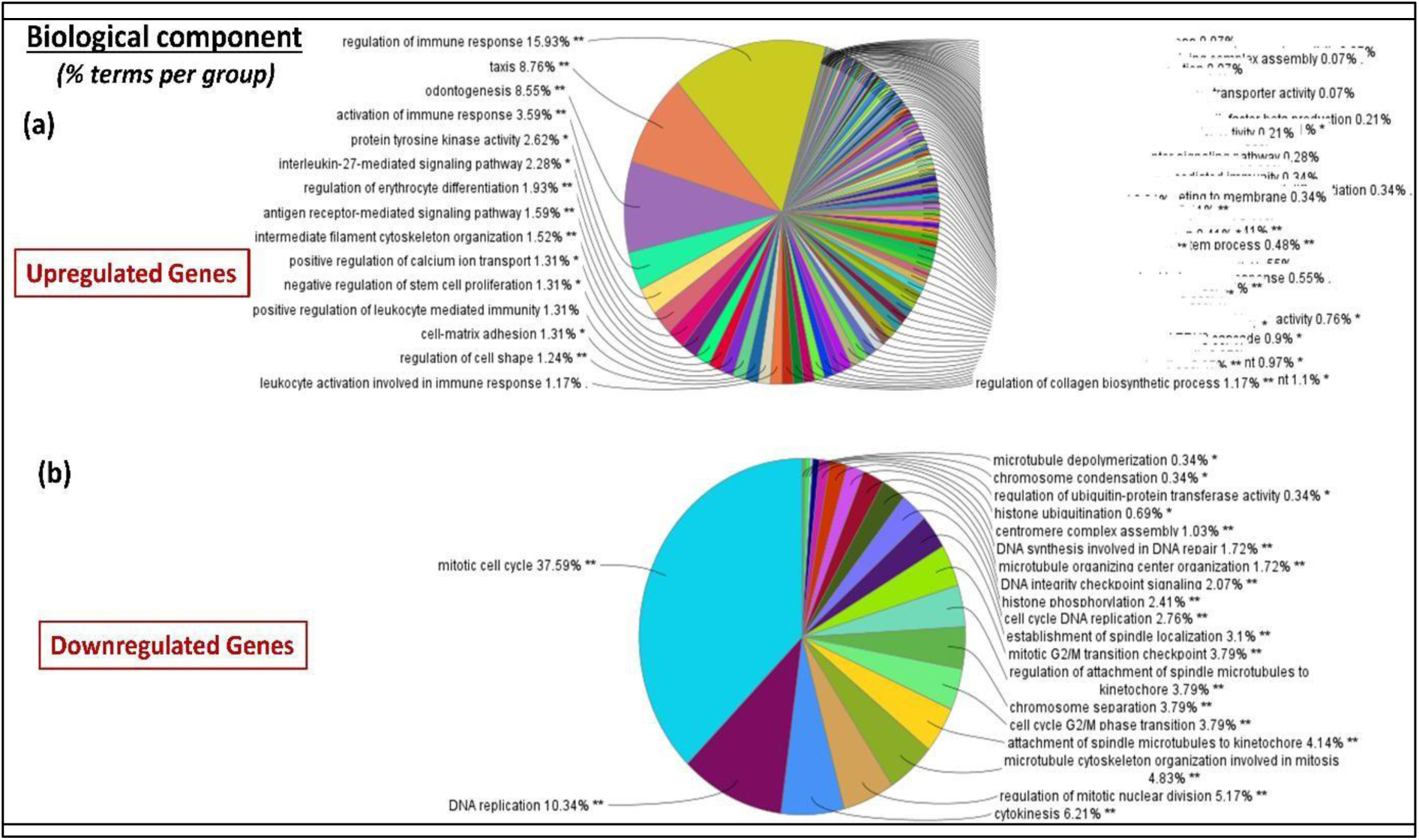
Biological Process (BP) of enrichment analysis: (a) Upregulated genes; (b) Downregulated genes.

*Downregulation:* On the other hand, the downregulated genes were mainly enriched in mitotic cell cycle, DNA replication, regulation of mitotic nuclear division, microtubule cytoskeleton organization involved in mitosis, cell cycle G2/M phase transition, establishment of spindle localization, cell cycle DNA replication and DNA synthesis involved in DNA repair **(Figure 4(b))**.

#### ***(iii)*** Cellular Component Group

*Upregulation:* The upregulated genes in the Cellular Component (CC) group were mainly enriched in both innate and acquired immunity entities, e.g. ficolin-1-rich granule (neutrophil), platelet alpha granule lumen releasing chemokines and cytokines, phagocytic vesicle, immune complement component C1 complex, etc. **(Figure 5 (a)).**

**Figure 5.**
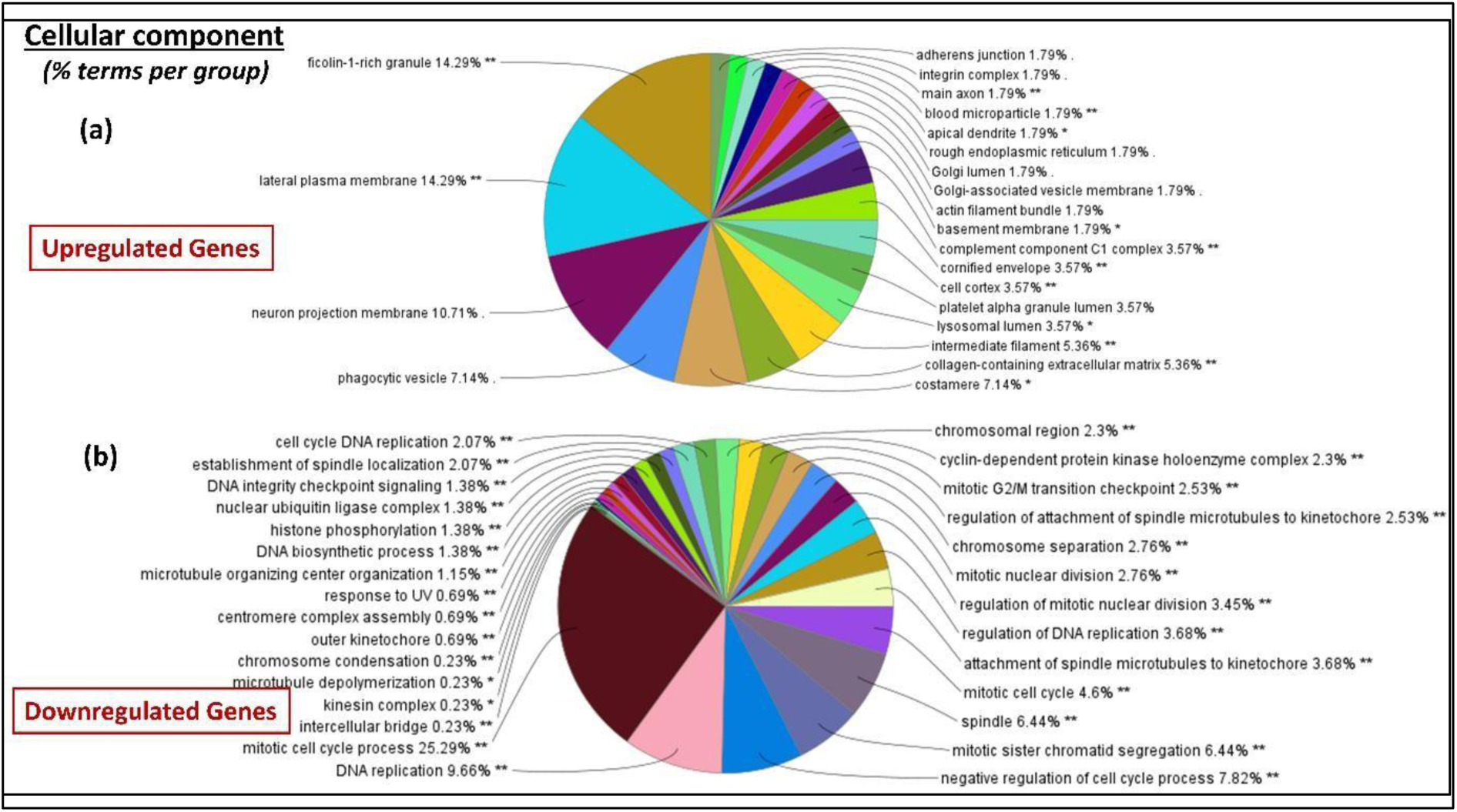
Cellular component (CC) of enrichment analysis: (a) Upregulated genes; (b) Downregulated genes.

*Downregulation:* In contrast, the downregulated genes were enriched in mitotic cell cycle process, regulation DNA replication, spindle, attachment of spindle microtubule to kinetochore, and chromosomes separation **(Figure 5 (b)).**

Regarding the Gene Ontology analysis we arrived at the following findings:

*Upregulation:* The majority of upregulated genes were involved in the activation and regulation of immune response as by activating neutrophils (the leukocytes that can act as the first line of defense of the immune system), maintaining the intercellular signaling, cytokine binding, etc. (**Figures 3(a), 4(a), 5(a)).**

*Downregulation:* In contrast, the downregulated genes were associated with the mitotic cell cycle process, spindle microtubule formation, DNA replication and cyclin-dependent protein serine/threonine kinase activity (**Figures 3(b), 4(b), 5(b)**) indeed, the inhibition of these genes is required for any malignant tumor in a host body to undergo the process of permanent regression.

Moreover, from the Biological pathway analysis, the upregulated genes were enriched in mesenchymal-to-epithelial transition, PAR1-mediated thrombin signaling events, thrombin/protease-activation receptor (PAR) pathway, integrin family cell surface interactions, and beta1 integrin cell surface interactions (**Figure 6(a)**). Activation of all these processes decreases the invasive ability of malignant cells by diminishing the mobility of these cells, by lessening the mobility of these cells and entrapping these cells. Also, the downregulated genes were mainly enriched in cell cycle, mitotic process, DNA replication, and G2/M checkpoints, G2/M DNA Damage checkpoints, and cell cycle checkpoints (**Figure 6(b)).** Indeed, these genes are strongly associated with the cell proliferation process.

**Figure 6.**
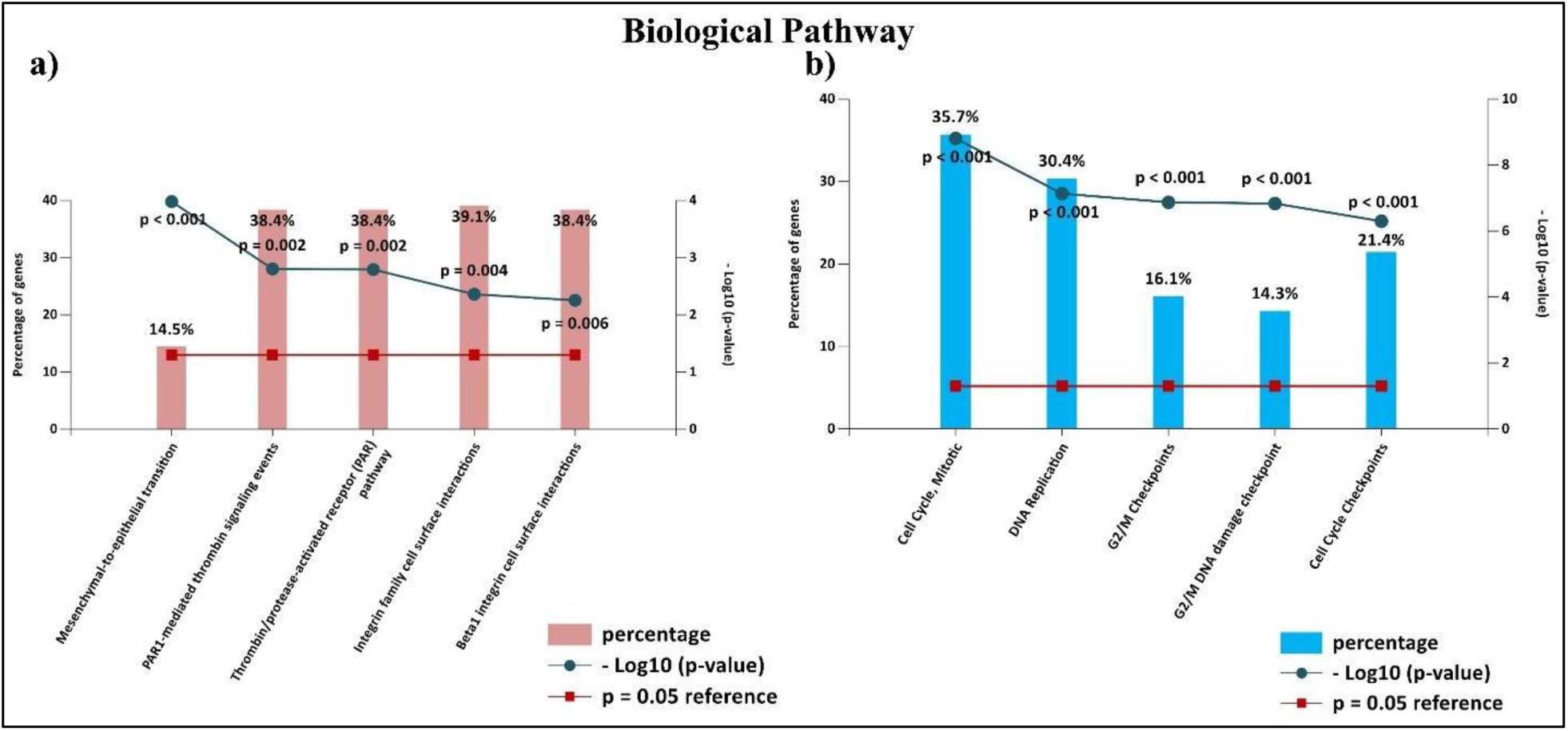
Significant Biological Pathway enrichment analysis of DEGs: (a) Upregulated DEGs; (b) Downregulated DEGs.

### 3.3 Identification of the Driving Genes Related to Regression of Melanoma Tumor

The top 10 hub genes were identified using cytohubba analysis for the first group of genes (**Table 1**), and we note therein that the majority of the genes are associated with DNA replication and its associated processes as cell division or mitosis/cytokinesis, and the genes are strongly downregulated in spontaneous regression process as the tabulated expression values show (i.e., downregulation of these genes indicate DNA interference or blockage effect during cell division). Further, we identified 17 common genes by taking commonality between microarray data, NCBI database genes, and gene card genes (**Table 2**) (**Supplementary Figure S2**). Hence, in order to explore alternate therapeutic strategies, we considered the mechanism of the spontaneous regression occurring in tumors, at the genetic as well as molecular level, therefore we started investigating the top 10 hub genes (Table 1) as well as the 17 common genes mentioned in Table 2. By extensive review of current information sources and documentation, we analyzed the pathways commonly associated with majority of these genes and found MAP kinase, JAK/STAT, PI3K/AKT as the major pathways to be dysregulated [49]. Thereafter, we explored these pathways and found RAS and RAF to be the protagonist proteins playing significant role in regulating these pathways [50].

**Table 1:** The top 10 hub genes with their fold change value pertinent to spontaneous regression of melanoma.

| Gene name | log FC value ($t_1$) | log FC value ($t_2$) | log FC value ($t_3$) | log FC value ($t_4$) |
| --- | --- | --- | --- | --- |
| TOP2A | -0.459517001 | -1.810469734 | -2.25435386 | -1.672954085 |
| KIF20A | -0.211683269 | -1.807779605 | -2.478033628 | -2.293056648 |
| KIF23 | 0.827249425 | 1.98418174 | 3.664080664 | 3.065167151 |
| CDK1 | -1.09722215 | -2.415204476 | -3.692431714 | -2.26071039 |
| CCNB1 | -0.636137771 | -1.936436546 | -2.738281085 | -2.068221461 |
| RAD51AP1 | -0.866296535 | -2.212584635 | -3.289193264 | -2.476289109 |
| BUB1B | -0.205789146 | -1.608319671 | -2.296878874 | -1.649891329 |
| CHEK1 | -0.510946563 | -1.22006972 | -1.519230504 | -1.091561664 |
| CCNA2 | -1.338388721 | -2.449128131 | -2.607963513 | -1.924035845 |
| KIF11 | -0.896395986 | -2.28949233 | -2.7775018 | -1.959737739 |

**Table 2:**
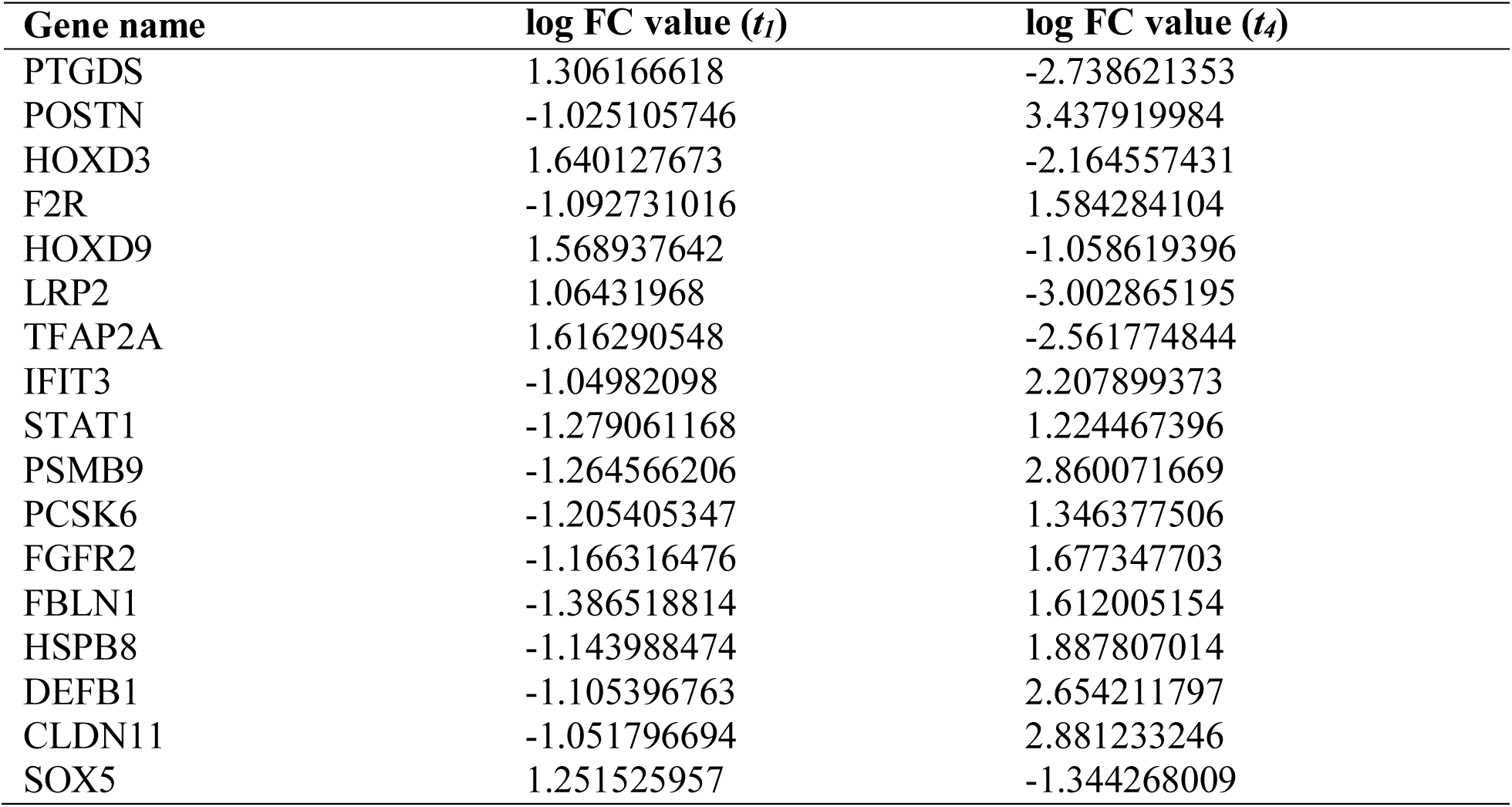
The 17 genes common between the microarray findings, NCBI database and Gene Card database with their fold change values. The genes have opposite signs in the expression values at time t_1_ (progression phase) and t_4_ (regression phase).

Further, investigating the downstream proteins and pathways associated, we found Cyclin D getting activated which ultimately leads to cell cycle progression/tumor progression **(Figure 7)**. Therefore, we decided to probe BRAF and NRAS, as these proteins are the initiators of the entire network of pathways and cell-cycle progression **(Figure 1(b))**. Both the MAPK/ERK (mitogen-activated protein kinase/extracellular signal-regulated kinase signaling pathway) and the PI3K/Akt (lipid kinase phoshoinositide-3-kinase signaling pathway) play a critical role in the transmission of cell signals via transduction systems (ligands, transmembrane receptors, and cytoplasmic secondary messengers) to the cell nucleus, where they affect the expression of genes that control crucial biological functions like cell growth, proliferation, differentiation and apoptosis [51,52]. The enrichment analysis of the total 27 driving genes also shows that the significant biological pathway was enriched in: (i) Chk1/Chk2 (cds1) - mediated inactivation of cyclin B: Cdk1complex, (ii) Cell cycle, Mitotic process, (iii) Cyclin a/B1 associated event during G2/M transition, (iv) Cell cycle checkpoints and (v) phosphorylation of protein involved in the G2/M transition by cyclin A: Cdc2 complexes. **Figure 8** shows the enrichment analysis. The aforementioned information is consistent with the notion that abnormalities in cell cycle checkpoints and cell growth regulators are the primary contributors to tumorigenesis [53,54].

**Figure 7.**
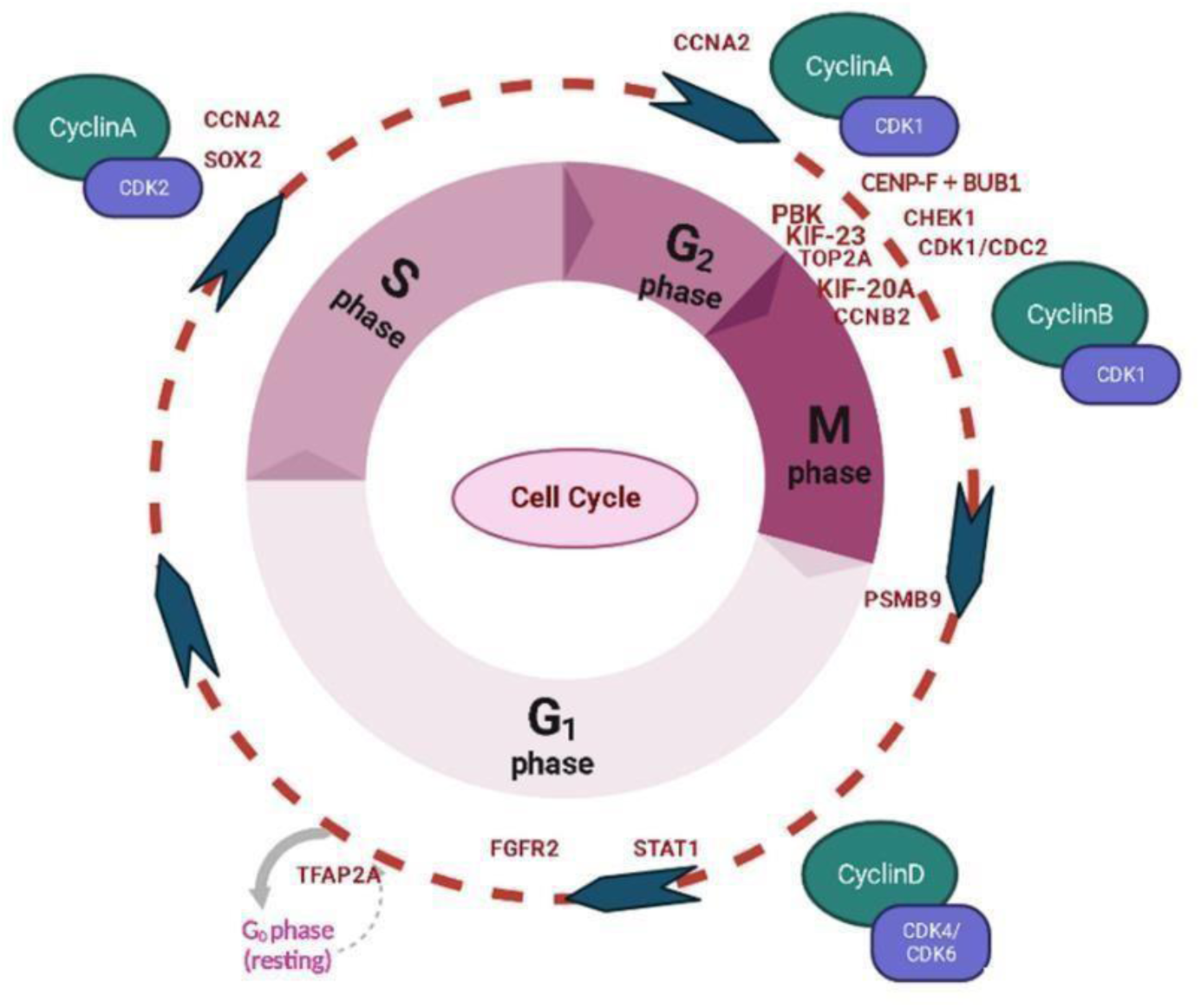
The cell cycle process under the genes that actuate the spontaneous regression phenomenon or permanent endogenous eradication of malignant melanoma tumor.

**Figure 8.**
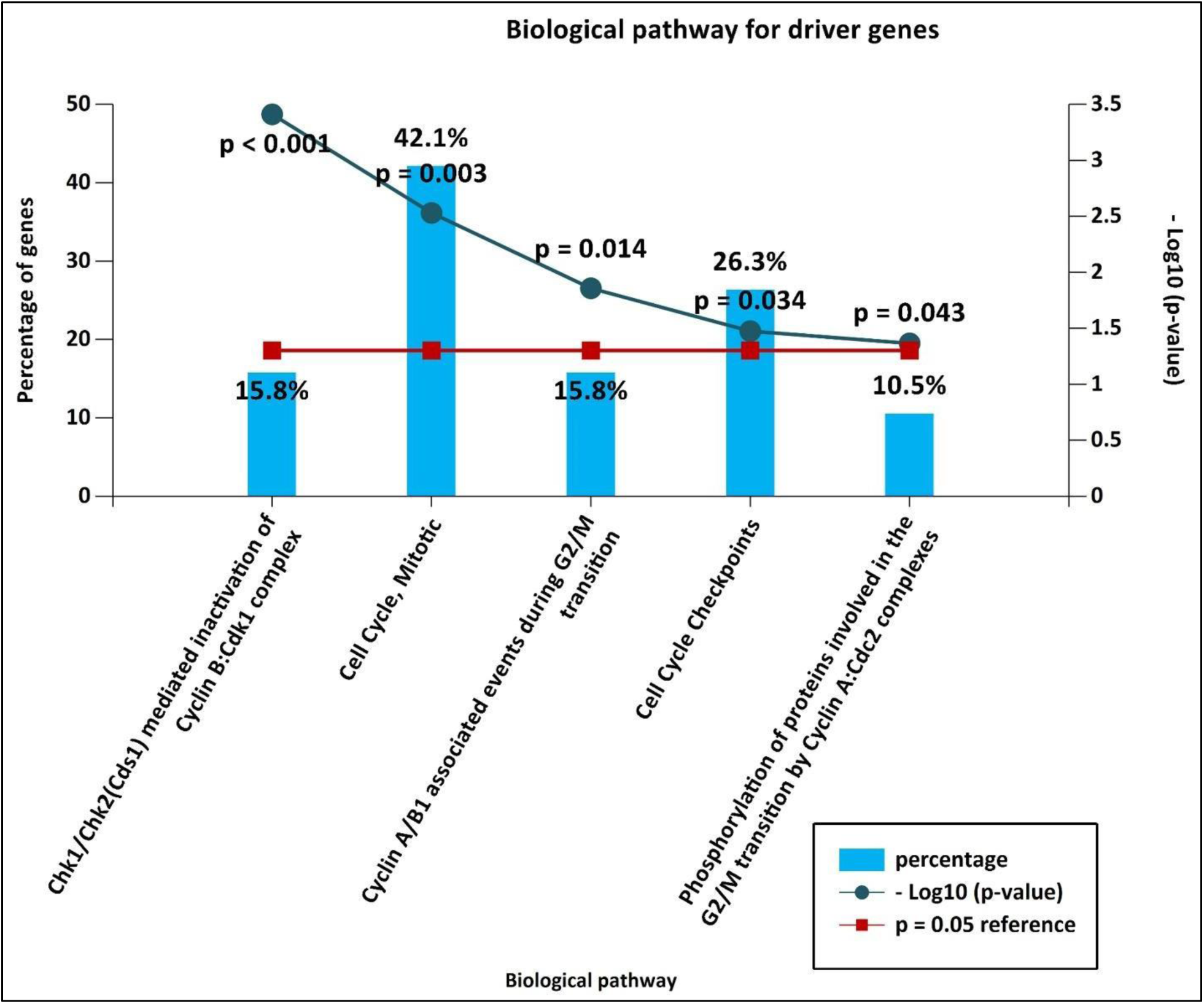
Enrichment analysis of significant Biological Pathway analysis of the driving genes pertinent to spontaneous regression of melanoma tumor.

### 3.4 Downstream signaling pathway of melanoma and its target protein

The mutually exclusive BRAF or NRAS mutations are the most frequent mutations that are present in melanoma. These mutations appear early in melanoma progression and continue throughout the disease’s progression [55]. Moreover, the NRAS and BRAF proteins help melanoma in progression through downstream activation of two signaling pathways: RAS-RAF-MEK-ERK pathway and PI3K-AKT pathway **(Figure 1(b))** which control cell growth, survival, and proliferation [23]. The pathways mediate the response of cells to mitotic extracellular stimuli. In the RAS-MAP pathway, the BRAF mutated gene acts on activated MEK, and then triggers the activation of the ERK downstream protein, which acts on DNA and enhances cell proliferation. Similarly, the mutated NRAS gene acts on PI3K and its activated PI3K pathway downstream. This activated phosphoinositide 3-kinase (PI3K) is typically the first step in the activation of the AKT pathway after exogenous growth factors have stimulated it. This is followed by increased production of the second messenger phosphatidylinositol-3,4,5-trisphosphate (PIP3), which can facilitate the translocation of AKT to the plasma membrane for its subsequent phosphorylation and activation. Since the phosphatase PTEN protein controls the intracellular level of PIP3, then PTEN’s functional deficit can cause the overexpression of PIP3 and induce AKT activation [56]. It is known that AKT3 is the most prevalent AKT isoform in melanoma. Furthermore, siRNA transfection directed against AKT3 or overexpression of PTEN are efficient ways to inhibit AKT3 activity and reduce the ability of melanoma cells to become tumorigenic [34]. As a result, the hyperactivation of the AKT pathway is a crucial oncogenic event for the development of melanoma.

### 3.5 Identification of Candidate Drugs

To screen out and select the drugs we considered three characteristics to select the drugs. Firstly, the drugs should interact with our proteins of interest viz., NRAS and BRAF, as analysed through DGIdb (Drug-gene Interaction database). It may be recollected that the drugs selected are Alpelisib, Obatoclax and Cetuximab (section 2.8.1). Thereby we found that Alpelisib has an interaction score of 0.14 with BRAF and 0.08 with NRAS (higher than other isoforms of RAS), and Obatoclax has an interaction score of 0.53 with NRAS and 0.25 with BRAF, while Cetuximab has an interaction score of 1.6 with BRAF and 0.4 with NRAS. Secondly, we have selected only those molecules which are FDA approved but have not been used for melanoma treatment. Thirdly, we shortlisted our final drugs with LibDock score. Relevant information on these candidate molecules are shown in **Supplementary Schema-1.** Since their molecular weight is below 500, they can cross blood-brain barrier, and hence may also be used in neurological melanoma, such as meningeal melanoma where meningeal melanocytes become malignant.

### 3.6 Molecular Docking with BRAF and NRAS proteins

The exploration of the binding of small molecules inside protein-ligand complexes as well as structure-based virtual screening have both made extensive use of molecular docking. Docking of our selected candidate drug molecules against the BRAF and NRAS gives several poses and the best pose for the individual docking complex was selected based on the best -CDOCKER score (lowest CDOCKER energy kcal/mol). In the case of protein-ligand docking, while in case of protein-protein docking the pose is selected based on the best E_RDock value (lowest one is the best). Results from the docking study are shown in **Table 3**. Further analysis of docking conformation within the binding pocket was done. 3D diagrams along with 2D interaction diagrams to visualize the interacting residues were also generated.

**Table 3.** The molecular docking score of the candidate drugs.

| Receptor Name | Ligand Name | CDOCKER Energy (kcal/mol) | CDOCKER Interaction Energy(kcal/mol) | LibDockScore |
| --- | --- | --- | --- | --- |
| BRAF | Alpselisib | -5.689 | -39.2787 | 90.2357 |
|  | Obatoclax | -15.6203 | -41.5001 | 90.9137 |
| NRAS | Alpselisib | -18.6548 | -66.7291 | 84.5067 |
|  | Obatoclax | -14.4835 | -50.8704 | 123.64 |
| Receptor | Antibody | ZDock Score | Z Rank Score | E_RDock (kcal/mol) |
| BRAF | Cetuximab | 17.24 | 13.172 | -12.2086 |
| NRAS | Cetuximab | 15.84 | -47.903 | -10.8281 |

#### Interaction study

The 3D representation of intermolecular interactions for both the protein complexes (BRAF and NRAS) with Alpelisib and Obatoclax drugs are given in **Figure 9 (a, b)** which have been constructed using Biovia Discovery Studio. Additionally, the 2D interactions are shown for the individual drugs, with receptors which help to visualize the types of interactions between the receptor residues and ligands like, Hydrogen bonds (imparting maximum binding strength), pi-cation bonds, pi-sigma bonds, vander waals forces, alkyl linkages, etc. enlisting the amino-acids involved in the interactions and thus stabilization of the interaction between the drug and the protein. The amino-acids colored in green show the hydrogen bond interactions, those in purple are the amino-acids interacting through pi-sigma bonds, orange colored amino-acids interact through pi-cation bonds and the amino-acids in pink represent alkyl linkages. These different amino-acids enlisted constitute the motifs on the proteins (BRAF and NRAS respectively) which are directly interacting with the drugs respectively **(Figure 9).**

**Figure 9.**
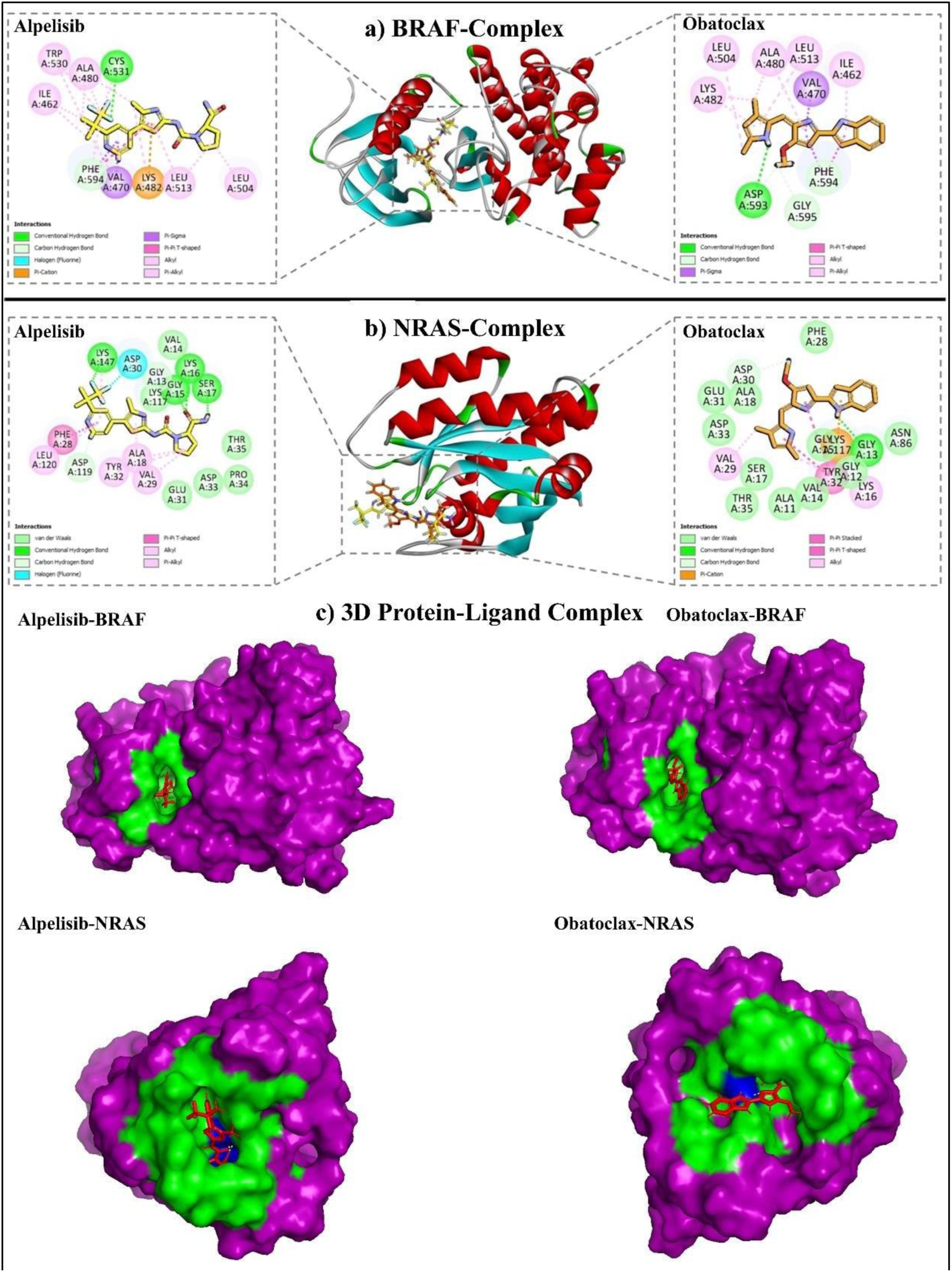
The 3D representation of the target receptors docked with ligands, illustrating the interacting residues in the 2D interaction diagram. (a) BRAF complex with Alpelisib and Obatoclax, (b) NRAS complex with Alpelisib and Obatoclax, (c) The 3D representation of Protein-Ligand complex.

As we can see in **Figure 9 (a)**, the number of bonds between BRAF and Alpelisib as well as Obatoclax are comparatively more than the number of bonds between NRAS and Alpelisib/ Obatoclax, as mentioned in **Figure 9 (b) [**also **Supplementary Table S1].** This infers that Alpelisib is a better interacting drug against BRAF and NRAS proteins as compared to Obatoclax. **Figure 9 (c)** illustrates the 3D complexes of protein-ligand structures, to show the surface representation of the protein. The solid surface of the protein (receptor) is represented in purple color, while the drug (ligand) is colored in red showing interactions marked with green color. Similarly, in **Figure 10**, the interactions of proteins (BRAF and NRAS) with Fab (Antigen-binding fragment) region of Cetuximab (monoclonal antibody) are demonstrated using Schrodinger’s Maestro software. Chain A and Chain B represent the heavy and light chains of the antibody (therefore dimeric structure). The amino-acids involved in the intermolecular interactions are displayed in stick representation with respective colors of protein and chains. Truncated line in black depict the hydrogen bond interactions with proteins, while those colored in green are showing the salt-bridges between the protein and the ligand (Antibody), and the dark green show pi-cation interactions.

**Figure 10.**
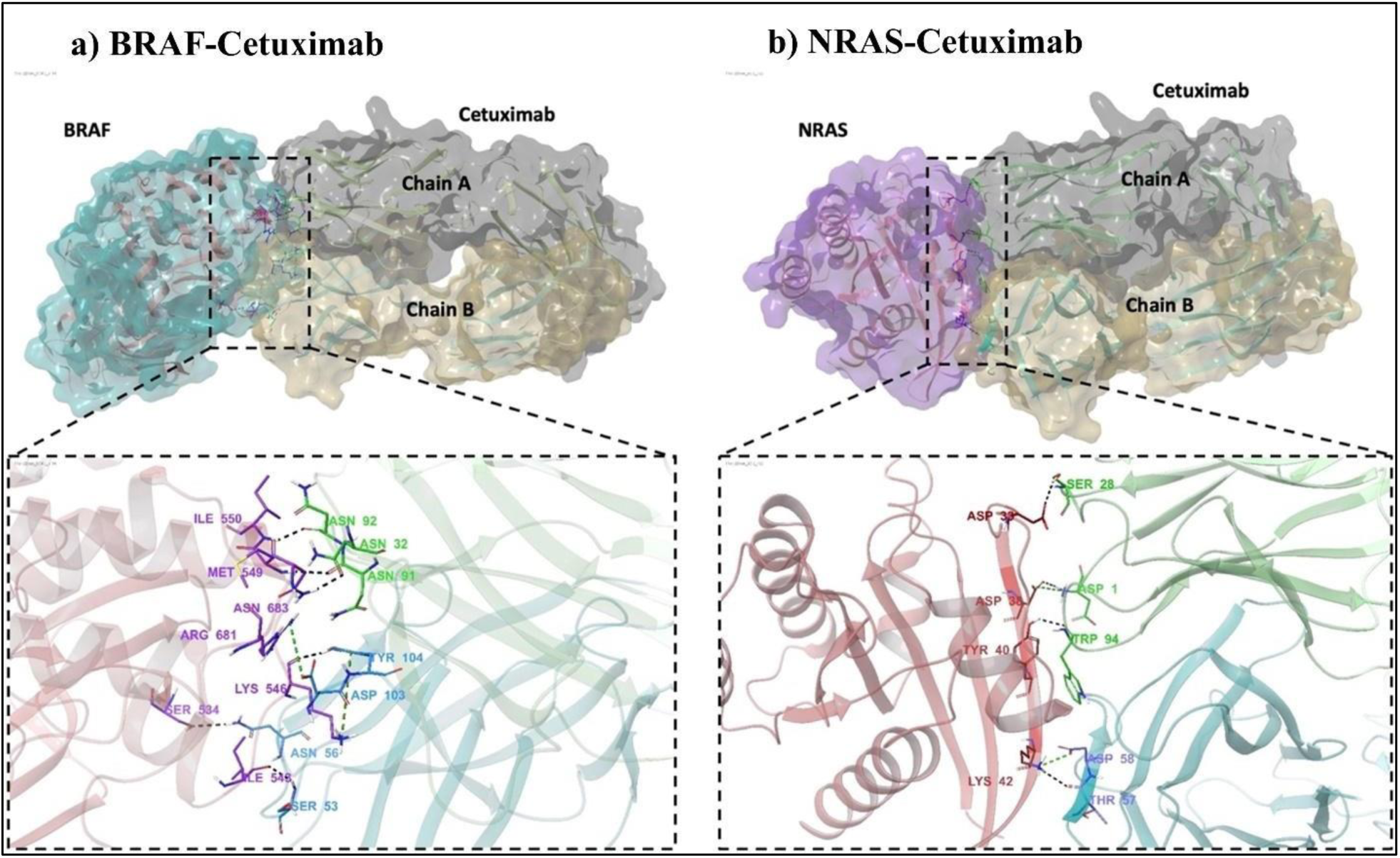
The 3D representation of the target receptors docked with Cetuximab, illustrating the interacting residues. (a) BRAF-Cetuximab docked complex, (b) NRAS-Cetuximab docked complex. Color scheme used for dashed boxes (lower panels) showing the interactions are as follows: hydrogen bond (black), salt-bridge (green), and pi-cation (dark green).

### 3.7 Molecular Dynamics Simulation Study

The investigation of molecular dynamics simulation enables us to delineate the atomic structure of the protein as well as its dynamic and structural characteristic changes occurring in real time [57]. The stability effect of our candidate molecules and monoclonal antibody were analyzed and studied for 100 ns, for each of the NRAS and BRAF proteins. This analysis was done based on RMSD, RMSF, radius of gyration, and intermolecular interaction (hydrogen bonds, salt bridges, pi-cation and pi-pi stacking).

In order to analyse protein stability, RMSD analysis was used to examine the conformational changes of protein atoms in relation to the reference frame. As it can be seen from the graphs, the conformation stability for Alpelisib shows that the ligand stabilizes after approximately 30 ns with each of the BRAF (5C9C) and NRAS (6ZIZ) respectively (**Figure 11 (a, b)**). Further, in the case of Obatoclax, there is stabilization with BRAF after approximately 30 ns nevertheless, there is a rigorous deviation of the ligand with NRAS **(Figure 11 (a, b))**. For Cetuximab, the deviation of the ligand is observed to be the same as the protein within the range of 2Å to 3.5Å (**Figure 11 (a)**) and 2Å to 3.8Å (**Figure 11 (b)**). These findings again demonstrate the stabilization of protein-ligand complex conformation.

**Figure 11.**
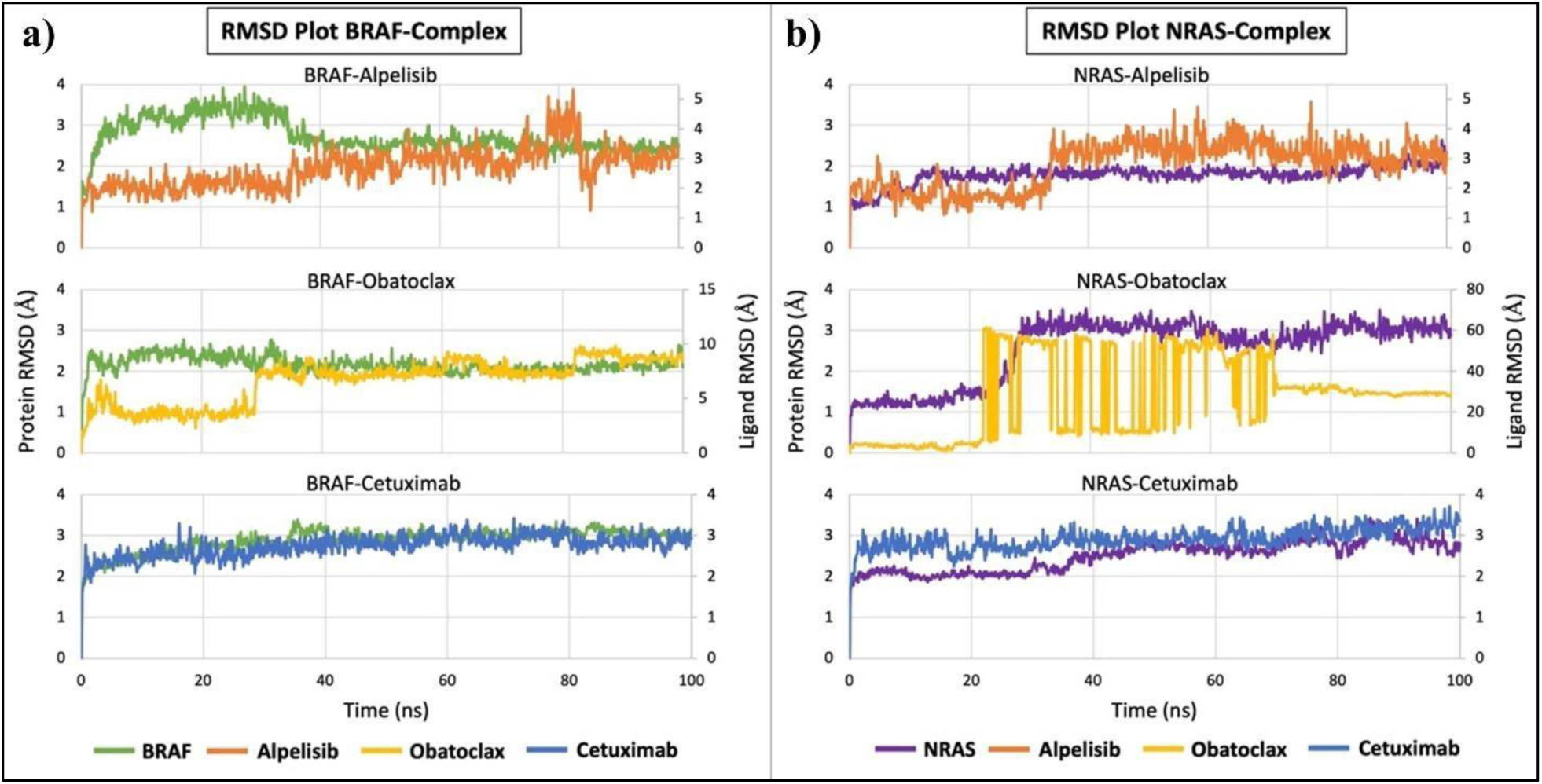
The RMSD analysis of protein-ligand complexes throughout the simulation trajectories. The RMSD of the protein is plotted on the left Y-axis, and the RMSD of the ligand is plotted on the right Y-axis, while simulation time (ns) is on the X-axis. (a) The RMSD of BRAF protein along with three selected ligands (Alpelisib, Obatoclax, and Cetuximab). (b) RMSD of NRAS protein along with three selected ligands (Alpelisib, Obatoclax, and Cetuximab).

The RMSF analysis helps characterize the local fluctuation along the protein chain. RMSF enables us to study the contributions of individual amino acids if they are stable or fluctuating during the simulation. The terminal and loop regions of the protein are extremely mobile. Here, the amino acid fluctuations in each protein-BRAF and protein-NRAS bound with the drug have been studied which demonstrates the fluctuation of the protein residues (**Figure 12 (a, b)).** Thus, as represented by the graphs, it can be seen that Alpelisib bound BRAF and NRAS entities show less RMSF values or fluctuations as compared to the Obatoclax and Cetuximab bound proteins **(Figure 12 (a, b)).**

**Figure 12.**
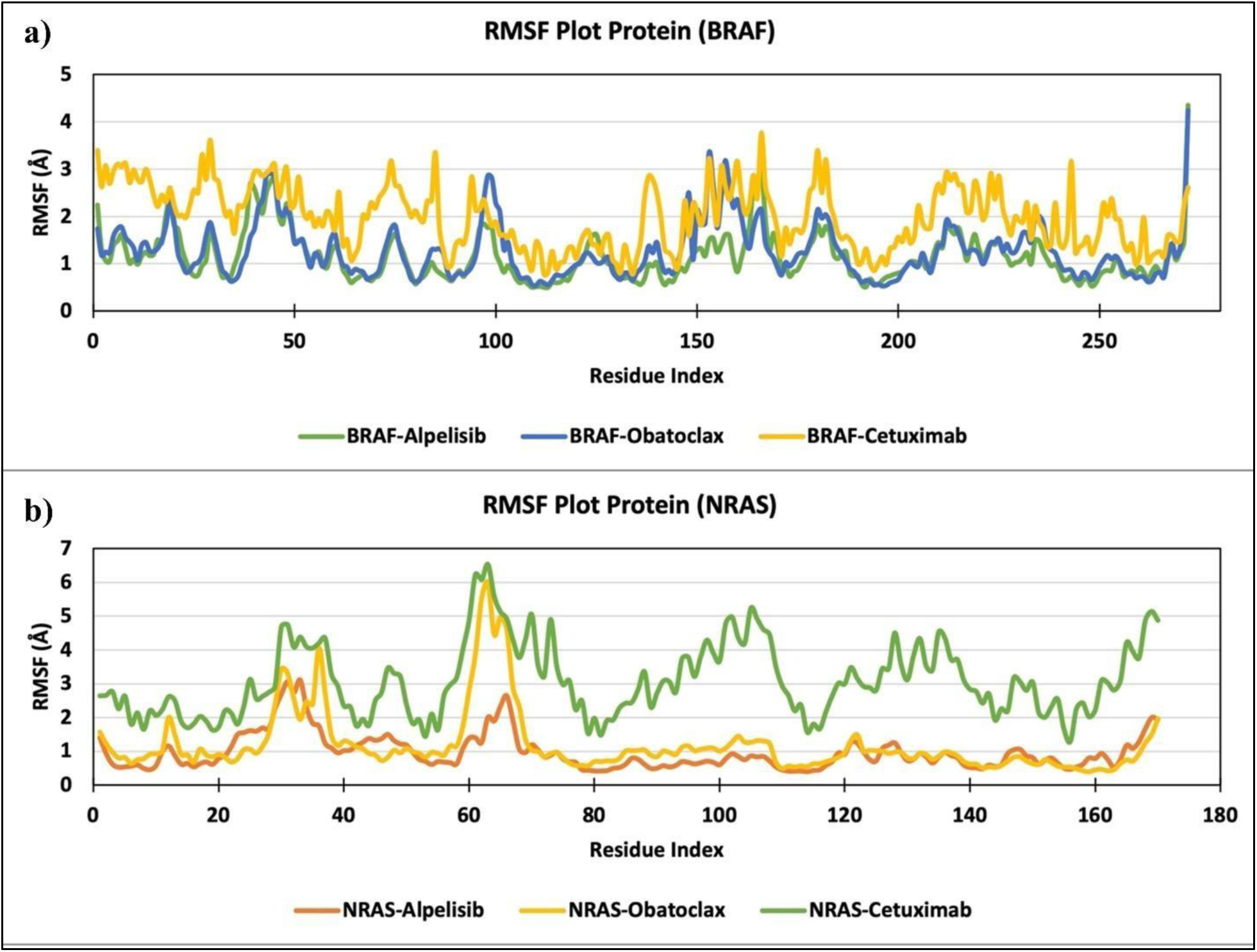
The RMSF analysis of protein C-alpha atoms for each residue throughout the simulation time, depicting the flexibility of the protein residues in the complexes. (a) RMSF plot of the BRAF complex; and (b) RMSF plot of the NRAS complex.

#### Intermolecular Interactions

Intermolecular interactions, such as hydrogen bonds (H-bonds) and salt bridges, enable proteins to maintain their conformation with the ligand and therefore, indicate the stability of the ligand-protein complex. Moreover, pi-cation and pi-pi stacking contribute to the binding of two molecules. We measured these interactions between the protein-ligand and protein-protein complexes in this study, to see the binding stability of complexes throughout the simulation trajectories of 100 ns each. It can be seen in the **Figure 13 (a, d)** that Alpelisib shows more hydrogen bond interaction with NRAS as compared to BRAF, while Obatoclax shows very minimal hydrogen bond interactions with either of the proteins **(Figure 13 (b, e)).** Additionally, in case of BRAF, Alpelisib and Obatoclax shows mostly pi-pi stacking type of interaction as compared to NRAS. We also observe that Cetuximab also shows continuous stable hydrogen bonding throughout the simulation with both BRAF and NRAS as shown in **Figure 13 (c, f)**.

**Figure 13.**
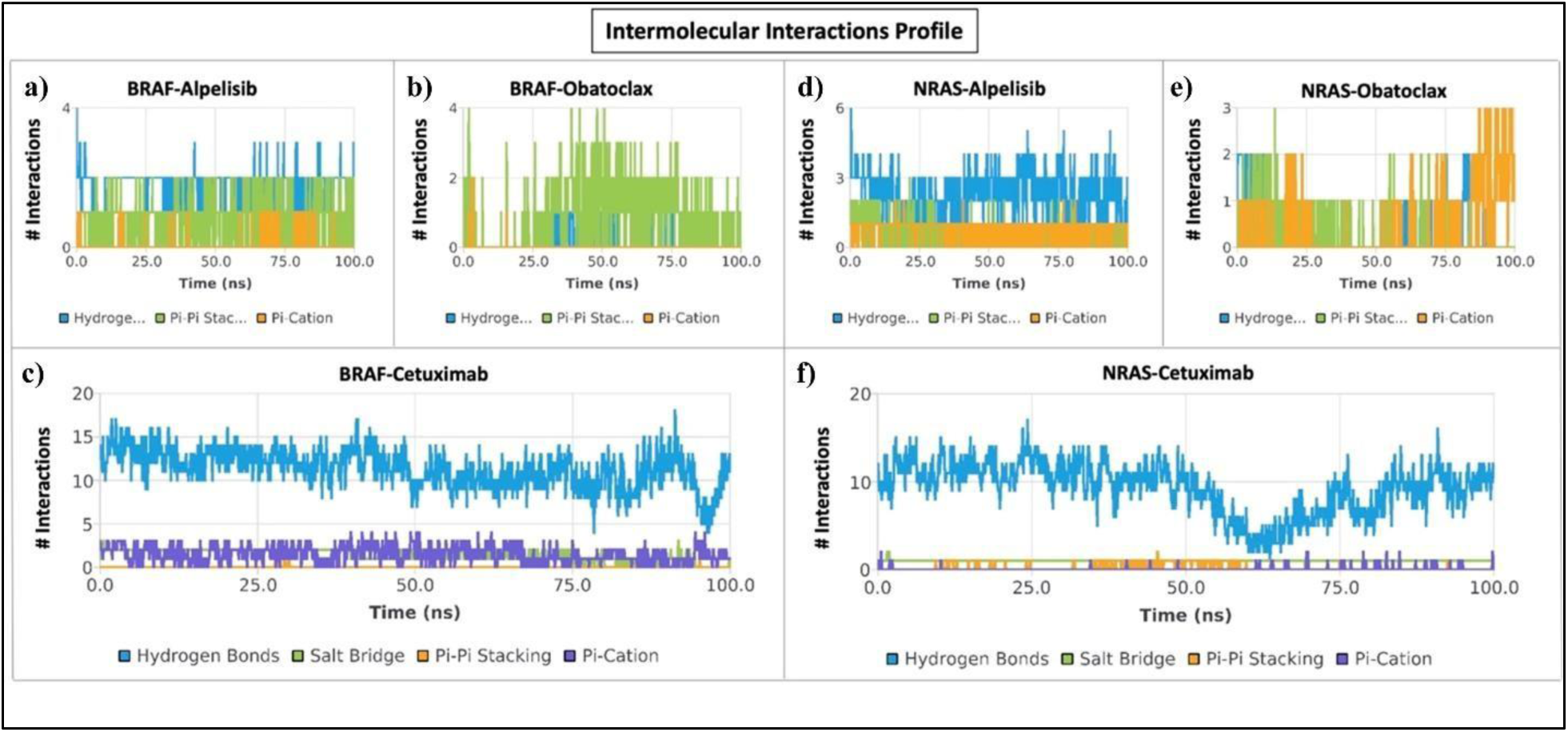
Intermolecular interactions between the protein and ligand as calculated during the simulation for estimating the stability of the complexes. The interaction profiles of the following complexes are depicted in the figure: (a) BRAF-Alpelisib, (b) BRAF-Obatoclax, (c) BRAF-Cetuximab, (d) NRAS-Alpelisib, (e) NRAS-Obatoclax, and (f) NRAS-Cetuximab.

Furthermore, the Radius of Gyration (Rg) measures the ‘compactness’ of the protein, which is calculated for the protein backbone and is equivalent to its principal moment of inertia. The radius of gyration (Rg) values for the compounds are as follows: Alpelisib exhibits a range of 19.1 Å to 20.5 Å for BRAF and 15.2 Å to 15.9 Å for NRAS (**Supplementary Figure S3 (a) and (d) respectively**). Obatoclax demonstrates a range of 19 Å to 19.9 Å for BRAF and 15.2 Å to 16.1 Å for NRAS (**Supplementary Figure S3 (b) and (e) respectively**). Cetuximab shows a range of 18.6 Å to 19.4 Å for BRAF and 15.2 Å to 16 Å for NRAS (**Supplementary Figure S3 (c) and (f) respectively**). Indeed, the Rg index determines the compactness of the protein during the simulation process. The overall radius of gyration analysis of BRAF and NRAS shows that these two proteins were not able to maintain their proper folding, and shows large changes during the simulation, except in case of the BRAF-Alpelisib complex.

#### Ligand-Protein Contacts

Further, the ligand-protein contacts have been analyzed and show the detailed interaction between the ligand atom and the protein residues **(Supplementary Figure S4**). Here, the proteins are BRAF and NRAF while the ligands are Alpelisib and Obatoclax. Gene regulation, immunoreaction, and signal transduction all depend on protein-ligand interactions. The efficacy of protein-ligand interactions is crucial for comprehending the dynamics of biological regulation and for developing and identifying novel therapeutic targets since it offers a theoretical framework for such investigations. In the chosen trajectory (0.00 through 100.00 ns), the interactions that take place more than 30.0% of the simulation time are displayed. In case of Alpelisib-BRAF complex, residues Leu513, Asp593, and Phe594 show contacts with more than 30% of simulation time, while in case of Alpelisib-NRAS, residues Gly15, Lys16, Ser17, Phe28, Glu31, Thr35, Lys117, and Ala146 interacted for more than 30% simulation time. This result indicates that Alpelisib shows better interaction with NRAS protein. Similarly, for the Obatoclax-BRAF complex, only residue Phe594 showed interaction, while in the case of the Obatoclax-NRAS complex, there was no interaction observed for more than 30% simulation time. This suggests that Alpelisib is a better interacting molecule as compared to Obatoclax against BRAF and NRAS proteins.

#### Ligand Torsion Profile

Moreover, for conformational evolution analysis, the ligand torsion profile was studied for every rotatable bond (RB) in the ligand, throughout the simulation trajectory (for 100 ns) (**Supplementary Figure S5 and Figure S6**). A torsion test is used to find out how a sample behaves when it is twisted, or when it is subjected to torsional stresses as a result of moments applied that cause shear stress around the axis. A two-dimensional schematic of a ligand with color-coded rotatable bonds is displayed in the top panel. A dial plot and bar plot of the same color is displayed with each rotatable bond torsion. Plots with dials, also referred to as radials, show how the torsion has conformed during the time duration. The bar plots provide an overview of the data on the dial plots by displaying the probability density of the torsion. Plotting the potential of the rotatable bond (by summing the potential of the corresponding torsions) is also possible if torsional potential information is available. The potential values are displayed in kcal/mol and are located on the left Y-axis of the chart. By examining the histogram and torsion potential correlations, one can understand the conformational strain the ligand experiences to retain a protein-bound conformation.

### 3.8 Binding Free Energy Calculations

**Table 4** presents the results of Prime MM/GBSA binding free energy calculations for protein-ligand complexes over the course of 100 ns of MD simulation. Each row in the table represents a different protein-ligand complex, and various parameters contributing to the binding free energy of each complex are measured and reported. For the BRAF_Alpelisib complex, the total binding free energy (ΔBind_Total) is calculated to be −63.504 ± 4.044 kcal/mol. Significant contributions to this energy include ΔBind_Coulomb (−9.282 ± 5.698 kcal/mol) and ΔBind_vdW (22.482 ± 5.276 kcal/mol), indicating the importance of electrostatic and van der Waals interactions in stabilizing the complex. In the case of BRAF_Obatoclax, the total binding free energy is determined to be −51.595 ± 6.815 kcal/mol. Notably, ΔBind_vdW (24.925 ± 3.061 kcal/mol) and ΔBind_Coulomb (−5.989 ± 2.288 kcal/mol) are the dominant contributors, underscoring the significance of van der Waals interactions in this complex.

**Table 4:**
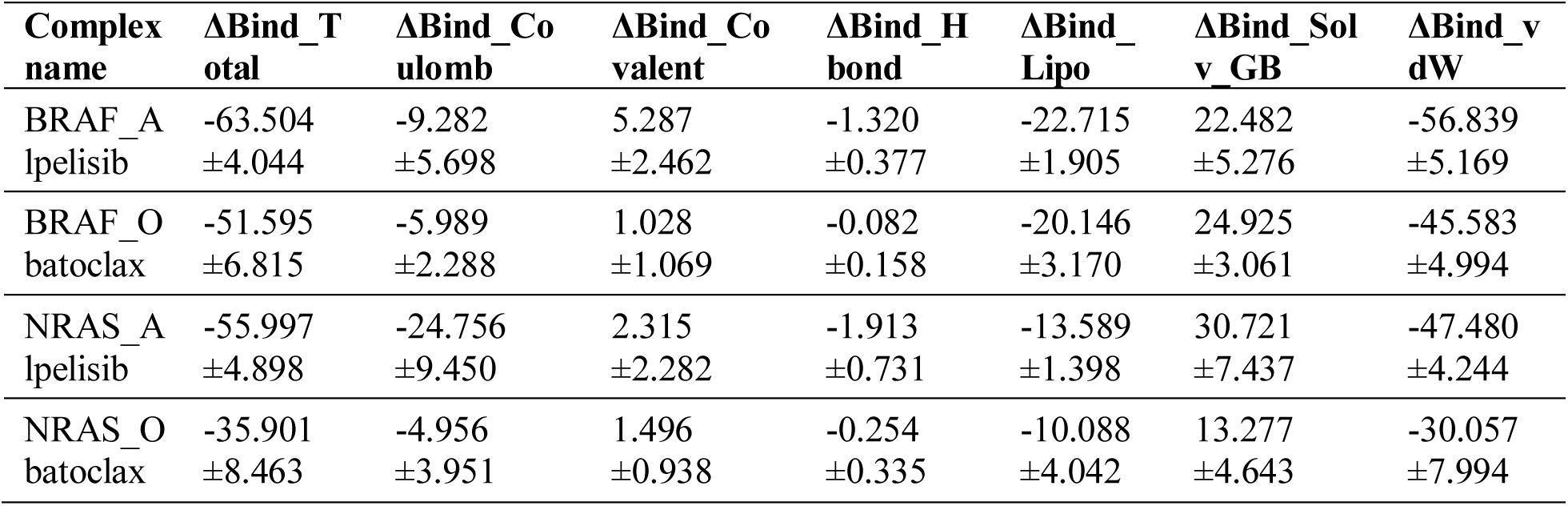
MM/GBSA Binding free energy calculation for the protein-ligand complexes during 100ns of MD simulation.

For NRAS_Alpelisib, the total binding free energy is found to be −55.997 ± 4.898 kcal/mol. Here, ΔBind_vdW (30.721 ± 7.437 kcal/mol) and ΔBind_Coulomb (−24.756 ± 9.450 kcal/mol) play crucial roles, suggesting strong van der Waals and electrostatic interactions in stabilizing the NRAS_Alpelisib complex. Finally, in the case of NRAS_Obatoclax, the total binding free energy is determined to be −35.901 ± 8.463 kcal/mol. ΔBind_vdW (13.277 ± 4.643 kcal/mol) and ΔBind_Coulomb (−4.956 ± 3.951 kcal/mol) are identified as major contributors, indicating the importance of van der Waals interactions in this complex as well.

## 4. Discussion

Numerous types of malignant lesions exhibit the episodic natural phenomena of spontaneous regression of cancer in which the tumor is totally eliminated permanently without toxicity effects. An in-depth investigation into this unusual natural occurrence may shed light on the feasibility of reproducing such a regression process on a clinical tumor in the patient context, without significant toxicity nor negative side effects. Here we developed a bioinformatics modelling, systematic analysis and therapeutic applicability regarding this regression process, with special reference to melanoma as an exemplar index case. The mechanism of the malignant growth process of melanoma has been the subject of numerous studies over the past few decades, nevertheless the incidence and death rates of melanoma are continuously rising globally. In contrast to earlier studies that only focused on several genes or a single platform or cohort, our work homed onto a spectrum of high-quality gene profile findings and datasets from several universal platforms in order to comprehensively investigate the driven genes and biological pathways involved in melanoma.

### 4.1 Oncological Targets and Drugs from Spontaneous Tumor Regression analysis

In our present study, we have investigated the spontaneously regressing melanoma microarray preparation and endeavored to explore the driver genes and biological pathways involved in melanoma regression. Finally, we have identified 344 upregulated and 160 downregulated DEGs. The enrichment analysis shows that the significantly downregulated genes were associated with the mitotic cell cycle process, DNA replication, G2/M checkpoints, and G2/M DNA damage checkpoint signaling pathways; these processes may play a crucial role in melanoma regression (**Figure 6(b)**). We have also identified that the 27 melanoma-driving genes are either involved in (i) cell cycle regulation and cell cycle division or (ii) signaling pathway activation, mainly as MAPK and PI3K/AKT signaling pathways. Furthermore, the enrichment analysis reveals that these 27 genes implicated in melanoma regression are enriched with regards to several key biological processes. These including: (i) cell cycle dynamics, (ii) mitotic processes, (iii) regulation of cell cycle checkpoints, particularly involving Chk1/Chk2 (cds1) mediated inactivation of cyclin B:Cdk1 complex, (iv) events associated with Cyclin A/B1 during G2/M transition, and (v) phosphorylation of proteins involved in the G2/M transition by cyclin A:Cdc2 complexes (**Figure 8**). Additionally, the cell cycle progression (**Figure 7**), highlighting specific genes or active sites, such as: (a) SOX2 genes interacting with cyclin D to regulate the cell cycle [58], (b) downregulation of FGFR2 gene leading to cell cycle arrest at the G1 phase [59], (c) interaction of STAT1 gene with cyclin D1/Cdk4 resulting in G1 phase arrest [60], (d) PSMB9 gene influencing the M/G1 transition phase [61], (e) TFAP2A affecting the resting phase of the cell cycle [62], and so forth.

Our investigation indicates that most of the DEGs were involved in either MAPK or PI3K/AKT signaling pathways or both. The RAS family proteins BRAF and NRAS, which are proteins associated with the oncogenes, are considerably frequent in malignant melanoma [63–65], and activates both those aforesaid signaling pathways. In our study, we have used computational analysis techniques like molecular docking and simulation dynamics to examine the inhibitory role of Alpelisib and Cetuximab on NRAS and BRAF genes, which are the most crucial genes in melanoma regression via inhibition of MAPK and PI3 kinase signaling pathways. Utilizing molecular dynamics studies, we have investigated the atomic/molecular level changes in the structural and physical aspects of the protein and even the individual amino acids.

### 4.2 Computational Biomolecular Analysis

Thereafter, we analyzed the changes in the behavioral pattern of the protein and stability of interactions of the ligand with each of the proteins by means of molecular dynamics simulation analysis using various parameters including RMSD, RMSF, Rg, and Protein-Ligand interactions (H-bond, Salt-bridge, pi-cation, and others). The conformational changes between the different candidate pharmacological agents interacting with our proteins have been visualized using a time-series analysis of the RMSD duration over the simulation. The plot thus generated for the protein-ligand complexes annotates the functional variations of the individual binding site residues, thereby depicting the changes in ligand binding behavior. Indeed, the RMSF calculations for each amino-acid residue of the protein annotate the measurements of local fluctuations throughout the simulation (100 ns). The radius of gyration (Rg) is an essential component in determining protein stability and protein folding patterns. A lower Rg value means that there has been proper folding or compactness of the protein, while, on the contrary, a higher Rg value indicates that aberrant protein unfolding has occurred.

By studying several interactions, such as H-bond and salt-bridges, the thermodynamic stability and folding pattern of the protein can be analyzed, since these bonds play an important role in the formation of secondary structure of proteins. Any changes in the bonding pattern could result in misfolding, thus indicating structural and thereafter functional impairments in the protein. This would further lead to alterations in the molecular pathways. The total binding free energy, as determined by MM/GBSA calculations, serves as a critical indicator of the strength and stability of protein-ligand complexes. In our study, the total binding free energies for the various protein-ligand complexes ranged from approximately −35.901 to −63.504 kcal/mol over the 100 ns MD simulation period. A key observation from the results is the variation in total binding free energies among different complexes. For instance, the BRAF_Alpelisib complex exhibited the highest total binding free energy of −63.504 kcal/mol, indicating a strong and stable interaction between the protein and ligand.

Conversely, the NRAS_Obatoclax complex displayed the lowest total binding free energy of −35.901 kcal/mol, suggesting a relatively weaker interaction compared to other complexes studied. These differences in total binding free energies can be attributed to several factors, including the chemical nature of the protein-ligand interface, the presence of specific binding motifs, and the overall structural complementarity between the protein and ligand. For example, complexes with higher total binding free energies may possess more favorable interactions, such as strong hydrogen bonding networks or extensive hydrophobic contacts, leading to increased stability. Furthermore, the total binding free energy serves as a valuable metric for evaluating the efficacy of potential drug candidates. Compounds with higher total binding free energies are more likely to form stable complexes with the target protein, thus increasing their potential as therapeutic agents. Reciprocally, compounds with lower total binding free energies may exhibit weaker binding affinities and may require further optimization to enhance their efficacy.

### 4.3 Translational perspective

Through the studies conducted, we have found that individually the candidate therapeutic molecules Alpelisib and Cetuximab, but not Obatoclax, show stable binding with both NRAS and BRAF. Thus, the two agents (Alpelisib and Cetuximab) can be re-purposed readily for inhibiting progression or for inducing regression of melanoma, since these drugs are already in clinical use for the treatment of several malignancies (but not melanoma) and are also FDA-approved. We have also found that the binding sites of Cetuximab and Alpelisib are not the same since Alpelisib is a small molecule (semi-synthetic chemical drug) binding to the active site of protein, whereas, Cetuximab is a monoclonal antibody binding to the variable region of the protein (antigen). Therefore, these two drugs can be used as a combination, which would allow for better efficiency and so the combination may need a lower dose for clinical efficacy. Moreover, the active pharmacophore moieties of Alpelisib and Cetuximab could be combined in a macromolecule which may be explored for a single dosing.

### 4.4 Overall clinical significance and implications of the study

We now furnish the incisive observations of our investigations:

a. The majority effect for actuating spontaneous melanoma regression is downregulation of DNA replication or mitosis promoting genes, i.e. this downregulation would parallel the effect of DNA interference or blockage (see sec. 3.3).
b. The next important facilitative factor for endogenous melanoma remission is activation of antitumor immune response, such as leucocyte activation and cytokine/chemokine signalling (sec. 3.2, items (i)-(ii)).
c. Reduction of metastasis ability of spontaneous melanoma remission is enabled by actuation of protease activation receptor 1 (PAR1) pathway (Fig. 6(a)).
d. This process of tumor eradication may be actuated conjointly by duplicating the two synergistic reciprocal coupled processes:

i. Enhancing the downregulation of spontaneous melanoma progression networks which can be enabled by agents as alpelisib and cetuximab (Fig. 9-10);
ii. Augmenting the upregulation of spontaneous melanoma regression networks that may be actuated by candidate molecules as teniposide and epriubicin (sec 1.2).
e. Though there are a vast plethora of confounding genes, pathways and drugs that are indicated in melanoma (8 major pathways, 9 major genes, 11 major drugs) (sec. 1.1), our analysis shows that only 2 genes (BRAF, NRAS), 2 pathways (Akt and Mapk) and 2 drugs (alpelisib and cetuximab) are main determinants of actuating permanent melanoma remission paralleling the spontaneous melanoma eradication process.
f. Our proposed therapeutic approach resembles the phenomenology of spontaneous cancer regression via exponentially decreasing trajectory which is the most optimal process for total tumor eradication, since there is minimal tissue damage and toxicity (sec. 2.1; Fig. 2(a)-(b)). It is this minimization aspect of toxicity and tissue hazard that characterizes the essence of spontaneous remission, namely there is no side-effects nor further tissue damage by malignant relapse.

As far as we know, this report is the first proposal of an innovative novel approach to clinical intervention in melanoma, orchestrating the reciprocal dynamics of both progression and regression processes, as clarified in item (iv) above (it may be mentioned that conventional interventions generally use the information and pathways of melanoma pathogenesis and progression dynamics that the physician attempts to retard, yet information on spontaneous melanoma regression is hardly clinically utilized).

## 5. Conclusion

We investigate the striking natural phenomenon of spontaneous remission of malignancy using computational biomolecular dynamics analysis, with applicability to malignant melanoma as an exemplar case. We highlight how this framework can be explored as a therapeutic approach of high novelty, since spontaneous cancer regression process is a very safe occurrence, and is never associated with toxicity nor recurrence, which are often precarious drawbacks of conventional antitumor therapies. As highlighted in the earlier subsection, it may be noted that the natural assured process of melanoma eradication (spontaneous remission) as well as our findings of the optimal drugs (alpesilib, cetuximab) shows that the DNA interference route is the primary effective route for melanoma extinction (though antitumor immunological activation is also necessary, but is subsidiary). This result that we obtained is unexpected as current clinical opinion regards melanoma immunotherapy as being more prospective that targeting DNA. Our insight of this anomalous but clinically significant issue would not have come to light unless we has investigated the natural spontaneous melanoma eradication process. Hence the seminal importance that further studies on endogenous melanoma remission behaviour be undertaken so as to have deeper therapeutic understanding for melanoma.

We identified possible candidate pharmacological molecules that could mimic the tumor regression process on the neoplastic lesion. Our current study indicates that the Ras-Raf-MEK-ERK and PI3K-PKT signaling pathways are seminally inhibited in spontaneous regression of malignant melanoma, and are complementarily activated in melanoma progression. This activation occurs either through NRAS or BRAF mutations, both of which arise early during melanoma pathogenesis and are preserved throughout tumor progression. Our present investigation points out that the candidate molecules Alpelisib and Cetuximab may potentially be novel repurposable drugs that can target both pathways together instead of targeting one at a time. We substantiated our approach using network pharmacoinformatics study and validated the feasibility of those candidate agents by drug docking simulation and molecular dynamics modelling, including the ligand-receptor study of binding affinity and binding free energy calculations, flexibility, stability, compactness, and the like.

Indeed, our proposed drugs may enable therapeutic duplication of the process of spontaneous regression phenomenon on malignant melanoma patients. The natural phenomenon of complete spontaneous cancer remission is a well-recorded protective process showing that various malignancies have the inherent ability to become eradicated and there is absence of toxic side effects. This tumor regression process is starkly unlike customary therapy-induced tumor regression, which often produce acute toxicity, treatment-refractory tumors and relapse of malignancy. This tissue homeostatic behavior of tumor regression is poorly understood, and having proper insight and comprehension of this process can have unique therapeutic applicability if this remission behavior could be duplicated on a malignant tumor in the patient. Our results highlight the seminal significance of computational systems biology analysis to reveal the pathways and targets that enable the permanent regression process in malignant melanoma as an index case study. Thereby we elucidate the substantially impactful implications for translational oncology, and we also identify the possible therapeutic molecules or candidate pharmacological agents that may duplicate a satisfactory tumor regression process on clinical subjects.

## Supporting Materials

Additional supporting documentation for this is available as Supplementary Materials.

## Data Availability Statement

a. Microarray Gene Expression Data is available on ArrayExpress platform (https://www.ebi.ac.uk/biostudies/arrayexpress/studies/E-MEXP-1152?query=E-%20MEXP-1152)
b. Molecular Dynamics analysis data is available on Mendeley Data platform having Reserved DOI: 10.17632/4xpff636jc.1

## Sponsorship

The sponsorship of La Dassault Systemes Fondation, France, is gratefully appreciated.

## Acknowledgements

PKR is thankful for support from C3i Hub Foundation, sponsored by Dept. of Science & Technology, Govt. of India. BK, AM and AS-1 are obliged for research fellowships from their respective institutions: [BK: I.I.T. (B.H.U.); AM & AS-1: Shiv Nadar Institution of Eminence, SNIoE]. AS-1 also acknowledges the Indian Council of Medical Research (ICMR), New Delhi, India, for providing a Senior Research Fellowship (Project ID: 2021-14351; File No.: BMI/11(100)/2022). KBL acknowledges SNIoE for the Post-Doctoral Fellowship. The support from Human Space Flight Program, Indian Space Research Organization (ISRO), Dept. of Space, Govt. of India, through the Respond scheme is sincerely appreciated. PKR acknowledges the facilitation of Shiv Nadar University for logistics support for this work.

## Author’s Contribution

*Bindu Kumari:* Conceptualization, Data curation, Formal analysis, Investigation, Methodology, Resources, Software, Validation, Visualization, Writing – original draft. *Arushi Misra:* Investigation; Methodology; Resources; Software; Validation; Visualization; Writing – original draft. *Ashish Shrivastava:* Investigation; Methodology; Resources; Software; Validation; Visualization; Writing - original draft. *Kiran Bharat Lokhande:* Investigation; Methodology; Resources; Software; Validation; Visualization; Writing - original draft. *Ashutosh Singh*: Resources; Software; Supervision; Writing – review & editing. *Prasun K. Roy:* Conceptualization, Formal analysis, Methodology, Resources; Software; Funding acquisition; Project administration, Supervision, Writing – review & editing.

