## Supplementary materials for "Spontaneous cancer regression as inversion of spontaneous cancer progression: Malignant melanoma remission as a case-study towards clinical applicability"

**Supplementary Schema, Table & Figures**

***Part 1: Supplementary Schema***

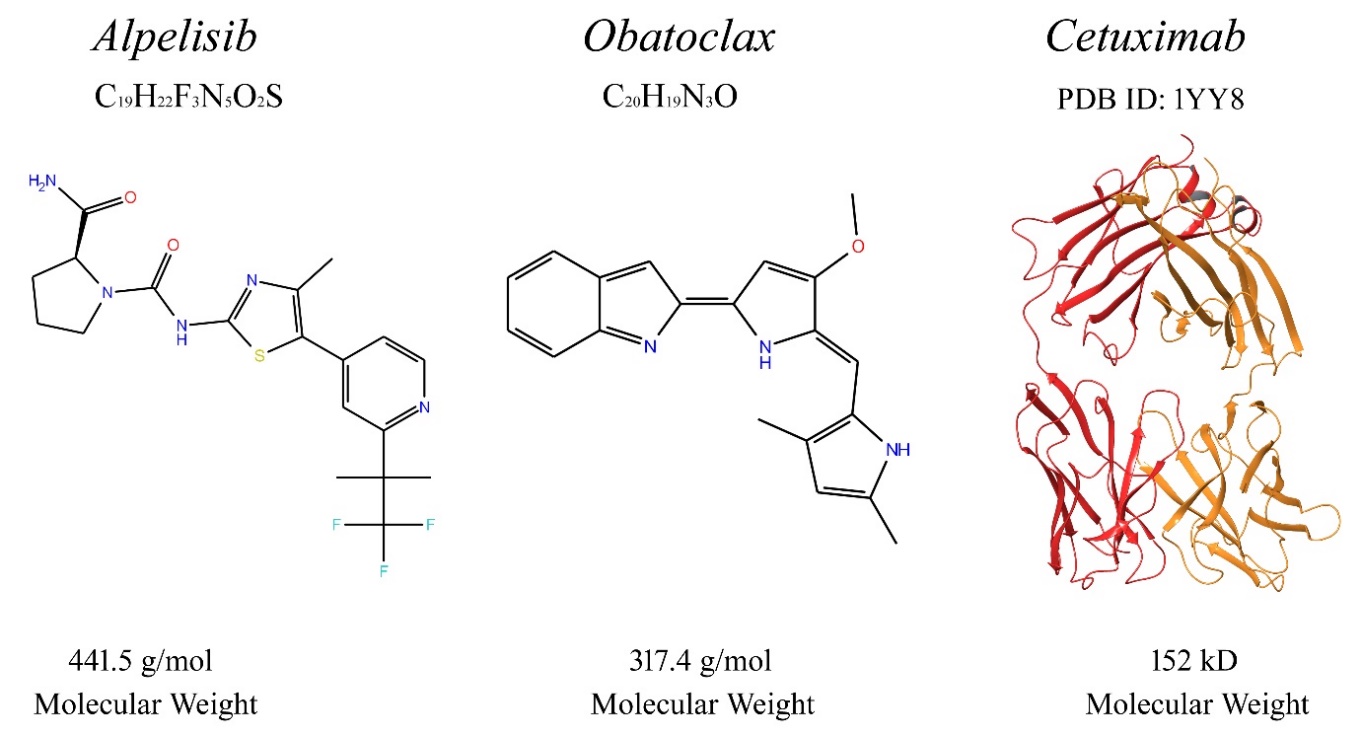
.

**Schema-1:** Details of the three candidate therapeutic molecules identified which may actuate a spontaneous regression-like process on melanoma tumor.

***Part 2: Supplementary Table***

| **S. No.** | **Complex Name** | **Residues** | | **Interaction Type** |
| --- | --- | --- | --- | --- |
| **Protein-Ligand Interaction** | | | | |
| 1. | BRAF-Alpelisib | Cys531 | | Conventional Hydrogen bond |
|  |  | Ala480, Trp530, Ile462, Leu513, Leu504 | | Alkyl linkage |
|  |  | Phe594 | | Carbon hydrogen bond |
|  |  | Val470 | | Pi-sigma bond |
|  |  | Lys482 | | Pi-cation bond |
| 2. | BRAF-Obatoclax | Asp593 | | Conventional Hydrogen bond |
|  |  | Gly595, Phe594 | | Carbon hydrogen bond |
|  |  | Val470 | | Pi-sigma bond |
|  |  | Ile462, Ala480, Lys482, Leu513, Leu504, | | Alkyl linkage |
| 3. | NRAS- Alpelisib | Gly15, Lys16, Ser17, Lys147 | | Conventional Hydrogen bond |
|  |  | Asp30 | | Halogen (Fluorine) bond |
|  |  | Phe28 | | Pi-Pi T-Shaped bond |
|  |  | Ala18, Val29, Leu120 | | Alkyl linkage |
| 4. | NRAS- Obatoclax | Gly13 | | Conventional Hydrogen bond |
|  |  | Lys117 | | Pi-Cation bond |
|  |  | Tyr32 | | Pi-Pi T-Shaped bond |
|  |  | Lys16, Val29 | | Alkyl linkage |
| **Protein-Protein Interaction** | | | | |
|  | | **Receptor** | **Antibody** |  |
| 5. | BRAF-Cetuximab | ASN92 | ILE550 | Hydrogen bond |
|  |  | ASN32/ASN91 | MET549 | Hydrogen bond |
|  |  | TYR104 | LYS546 | Hydrogen bond |
|  |  | SER53 | ILE548 | Hydrogen bond |
|  |  | ASP103 | ASN683/ARG681 | Salt-bridge |
|  |  | ASN56 | SER534 | Pi-Cation bond |
| 6. | NRAS-Cetuximab | SER28 | ASP33 | Hydrogen bond |
|  |  | TRP94 | TYR40 | Hydrogen bond |
|  |  | THR57 | LYS42 | Hydrogen bond |
|  |  | ASP1 | ASP38 | Pi-Cation bond |
|  |  | ASP58 | LYS42 | Salt-bridge |

**Table S1:** Intermolecular interaction analysis of ligands with receptor proteins (BRAF, NRAS).

***Part 3: Supplementary Figures***

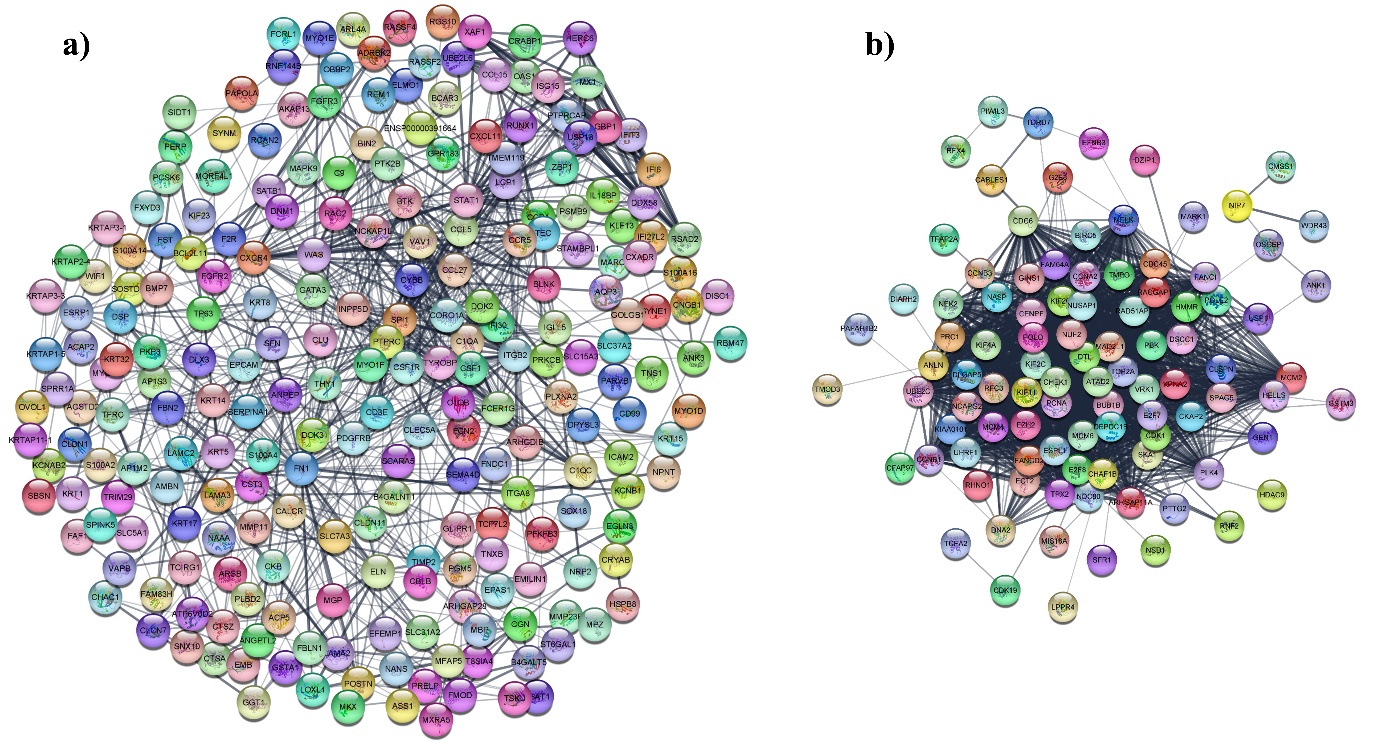

**Figure S1.** The Protein-Protein interaction (PPI) network: (a) Upregulated differentially expressed genes (DEGs); (b) Downregulated differentially expressed genes (DEGs).

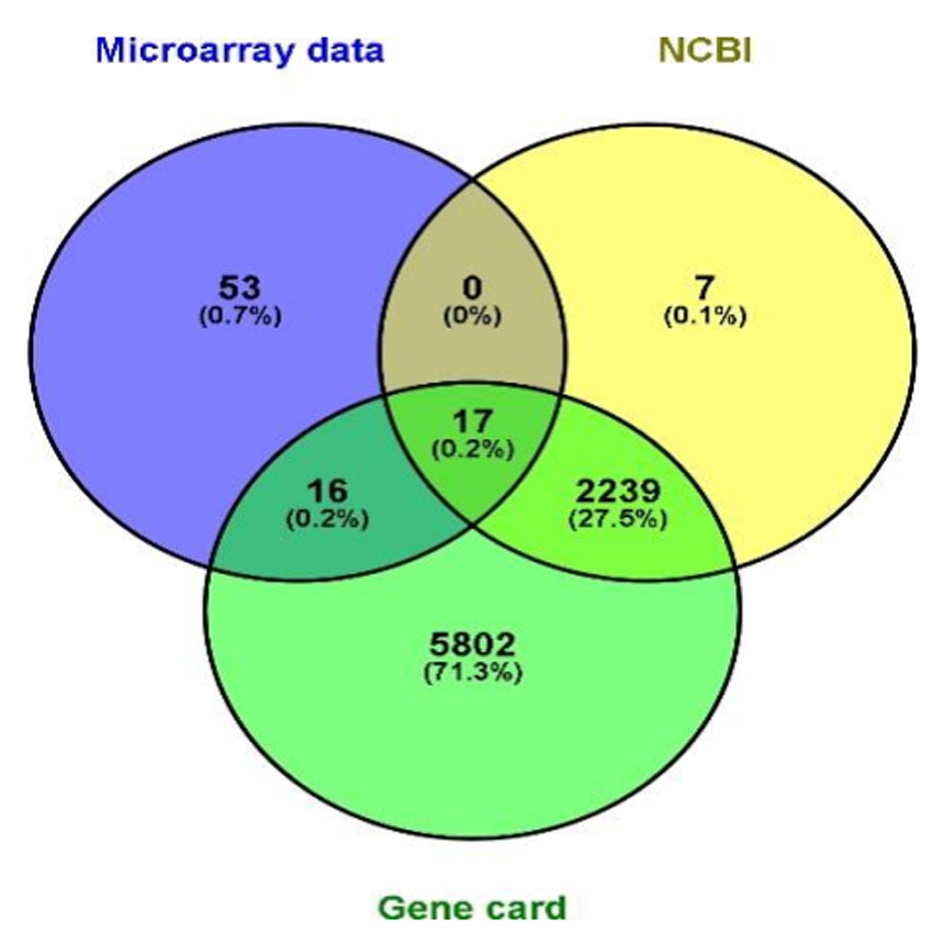

**Figure S2**. Venn diagram representing the common genes between the Microarray findings, the NCBI database and the Gene Card database.

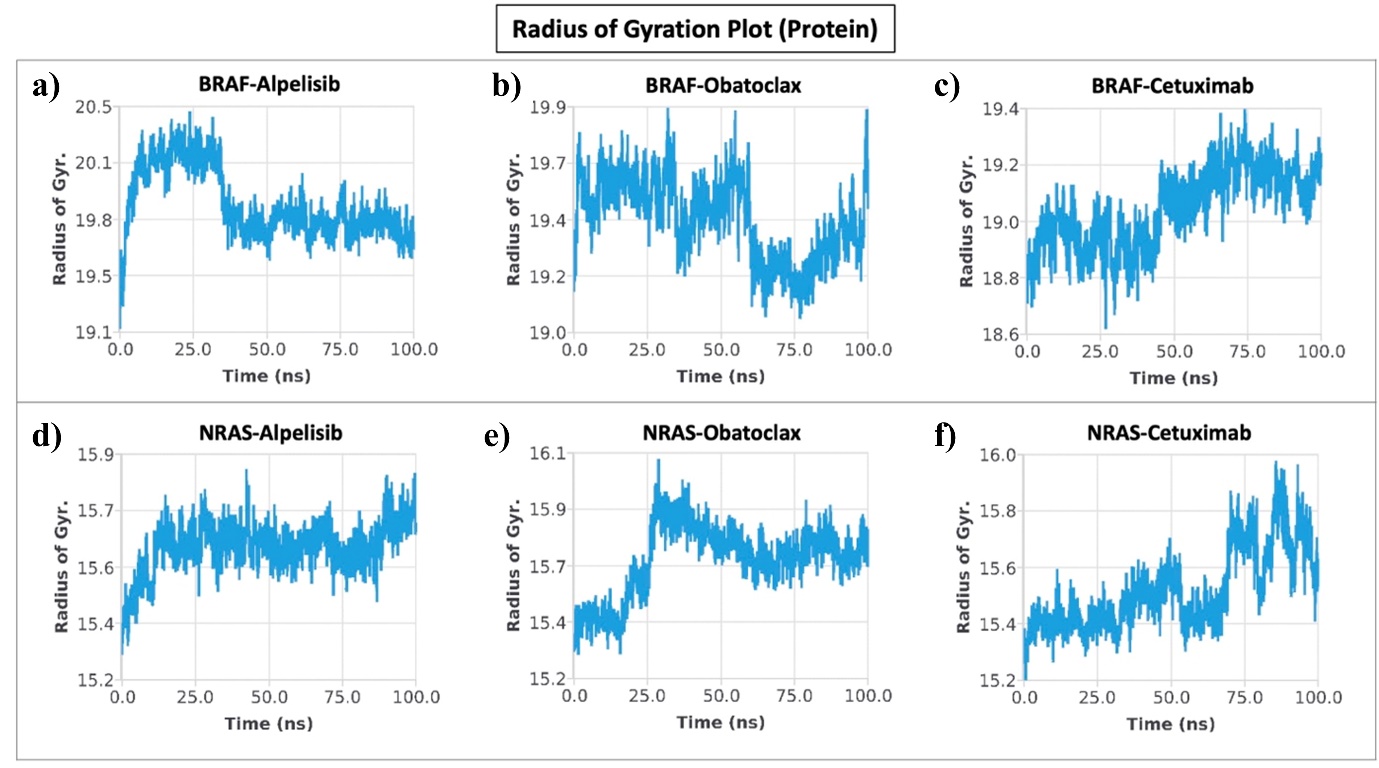

**Figure S3.** The radius of gyration plots of the protein-ligand complex, displaying the changes in protein compactness throughout the simulation. The radius of gyration of the protein is depicted for each complex: (a) BRAF-Alpelisib complex, (b) BRAF-Obatoclax complex, (c) BRAF-Cetuximab complex, (d) NRAS-Alpelisib complex, (e) NRAS- Obatoclax complex, and (f) NRAS-Cetuximab complex.

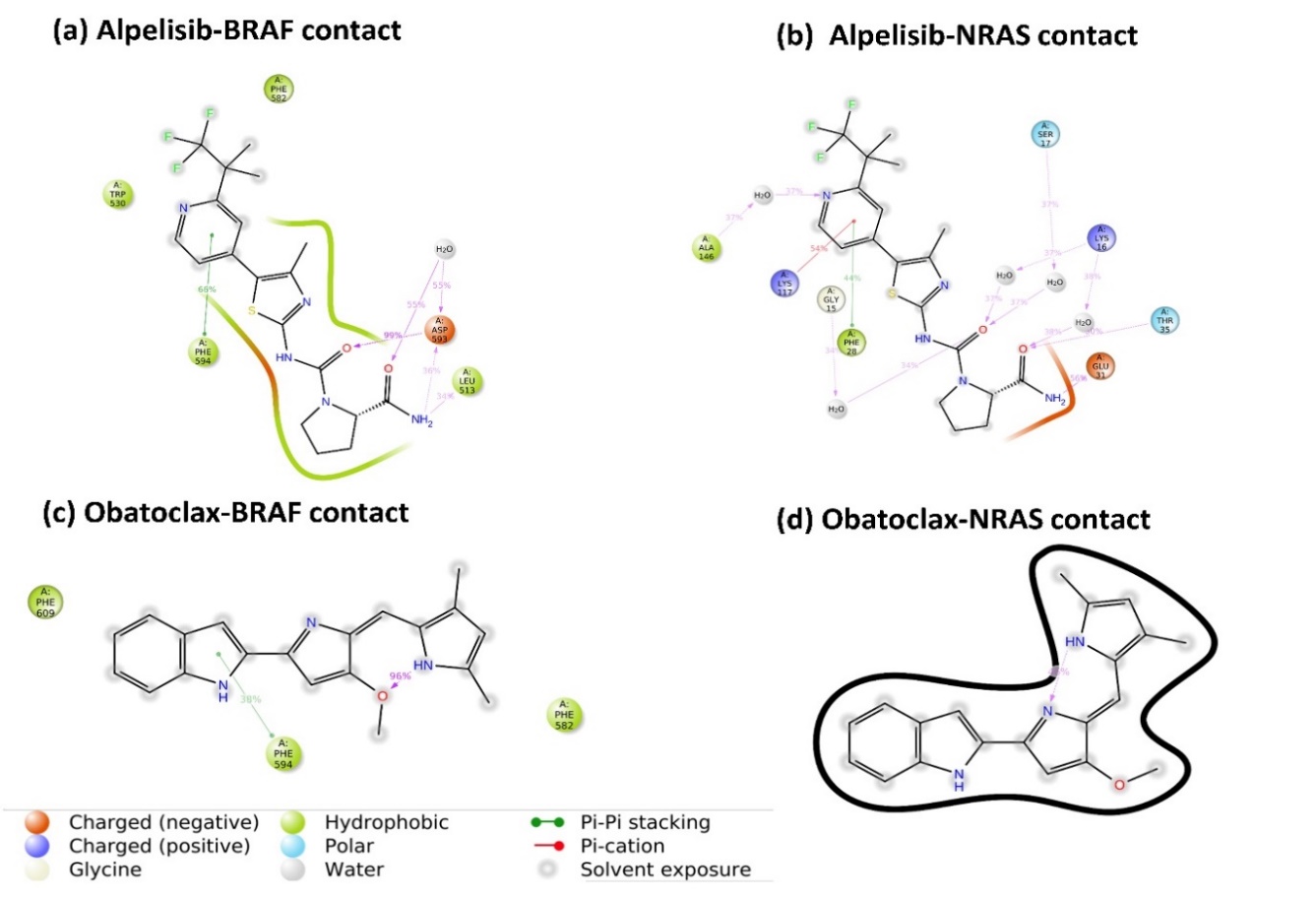

**Figure S4.** Ligand atom interaction with protein residues. Interactions that occurs more than 30 % during whole simulation time (i.e. for 100 ns) are shown here.

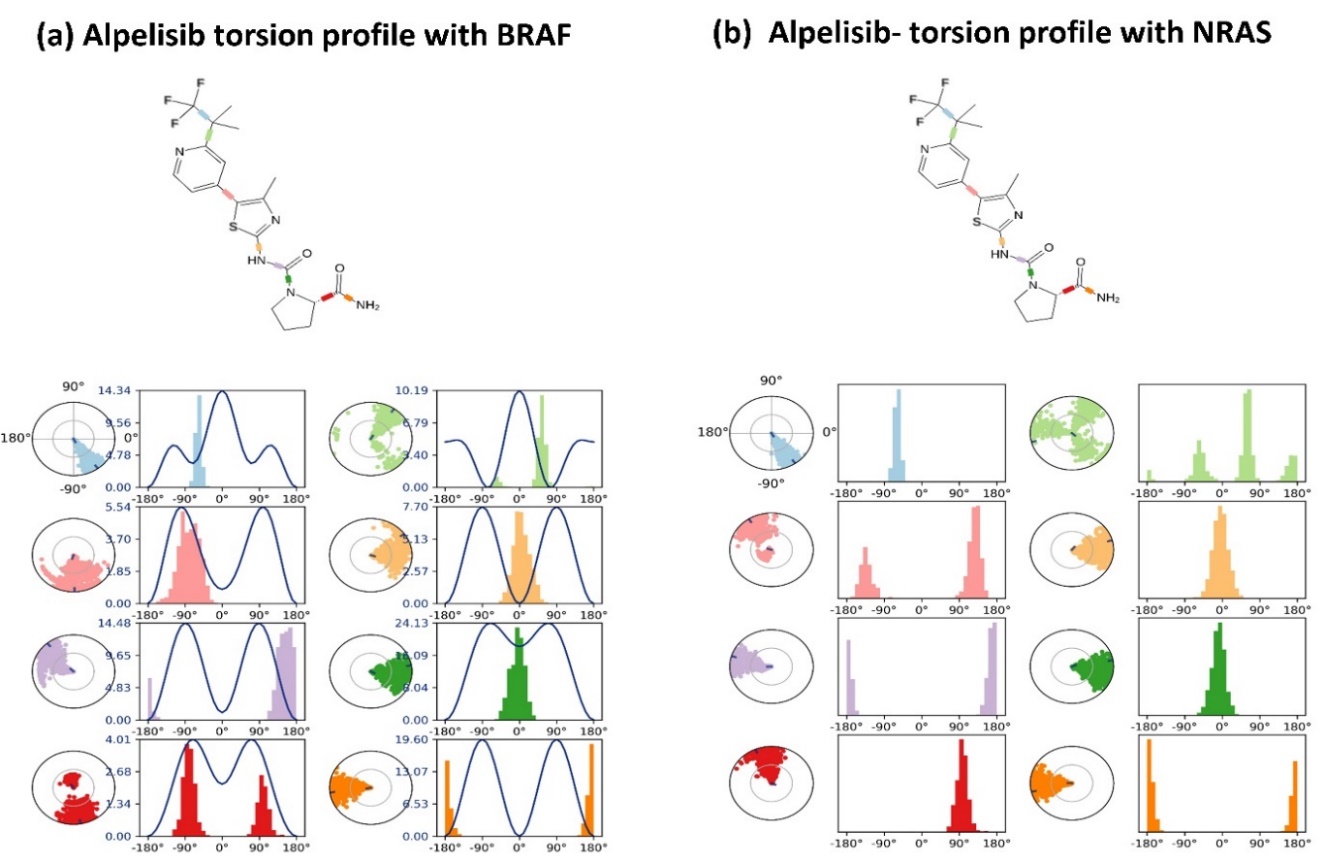

**Figure S5**. Ligand torsions plot, which gives conformational evolution of every rotatable bond throughout the simulation from 0 ns to 100 ns: (a) For Alpelisib with BRAF; (b) Alpelisib with NRAS; the upper panel shows the torsion and flexibility and the lower panel shows the torsion angles.

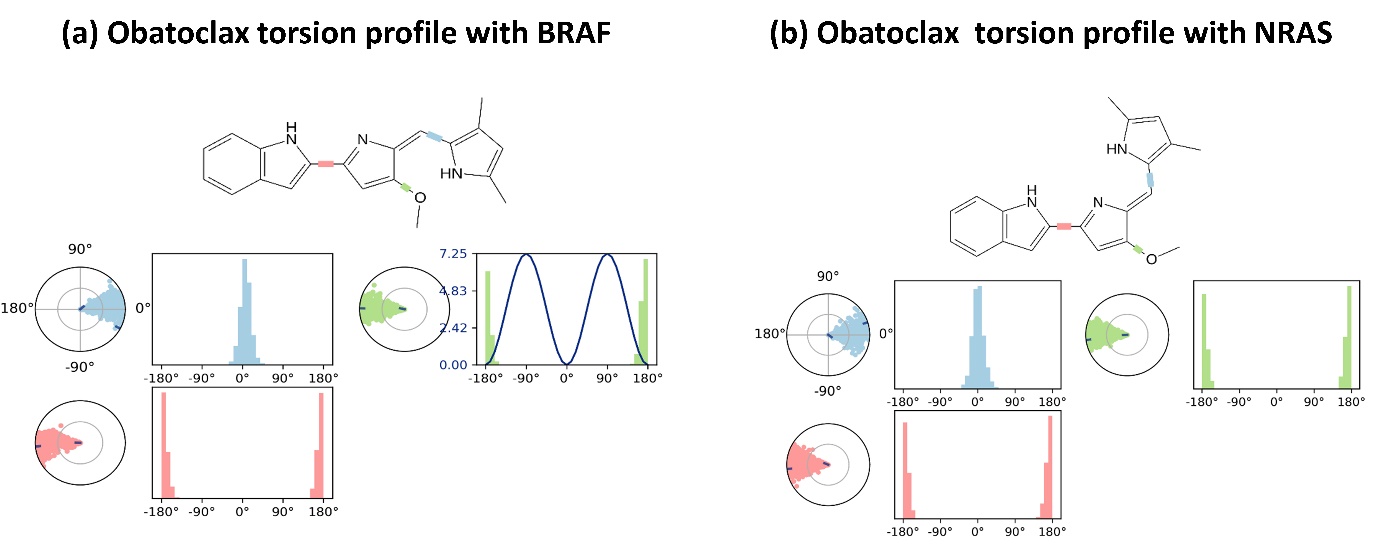

**Figure S6**. Ligand torsions plot, which gives conformational evolution of every rotatable bond throughout the simulation from 0 ns to 100 ns: (a) For Obatoclax with BRAF; (b) Obatoclax with NRAS; the upper panel shows the torsion and flexibility and the lower panel shows the torsion angles.
